# Enzyme Structure Correlates With Variant Effect Predictability

**DOI:** 10.1101/2023.09.25.559319

**Authors:** Floris van der Flier, David Estell, Sina Pricelius, Lydia Dankmeyer, Sander van Stigt Thans, Harm Mulder, Rei Otsuka, Frits Goedegebuur, Laurens Lammerts, Diego Staphorst, Aalt D.J. van Dijk, Dick de Ridder, Henning Redestig

## Abstract

Protein engineering increasingly relies on machine learning models to computationally pre-screen promising novel candidates. Although machine learning approaches have proven effective, their performance on prospective screening data leaves room for improvement; prediction accuracy can vary greatly from one protein variant to the next. So far, it is unclear what characterizes variants that are associated with large prediction error. In order to establish whether structural characteristics influence predictability, we created a combinatorial variant dataset for an enzyme, that can be partitioned into subsets of variants with mutations at positions exclusively belonging to a particular structural class. By training four different variant effect prediction (VEP) models on structurally partitioned subsets of our data, we found that predictability strongly depended on all four structural characteristics we tested; buriedness, number of contact residues, proximity to the active site and presence of secondary structure elements. These same dependencies were found in various single mutation enzyme variant datasets, with effect directions being specific to the assay. Most importantly, we found that these dependencies are highly alike for all four models we tested, indicating that there are specific structure and function determinants that are insufficiently accounted for by popular existing approaches. Overall, our findings suggest that significant improvements can be made to VEP models by exploring new inductive biases and by leveraging different data modalities of protein variants, and that stratified dataset design can highlight areas of improvement for machine learning guided protein engineering.

## 1 Introduction

Enzymes are nature’s catalysts, allowing reaction rates that are orders of magnitude higher than those of their non-biological counterparts (1). They are also versatile, catalyzing many different reactions that are relevant in addressing societal challenges, such as carbon capture (2) and plastic degradation (3). Enzymes are used in many commercially relevant applications, such as stain removal on fabric or dish surfaces, starch degradation for beer brewing or glucose production from plant waste. Protein engineering seeks to optimize the performance of enzymes in the context of an application through the introduction of mutations. These mutations are often selected by variant effect prediction (VEP), a field that is of interest not only to protein engineering, but also in medicine (4) and agriculture (5), where mutations induce changes in phenotype.

Machine learning models that predict the effect of mutations can guide protein engineering by computationally pre-screening protein variants in numbers that far exceed the capacity of experimental screens. Machine learning models used in protein engineering set themselves apart from other domains of VEP in that the properties/functions of interest are often non-native, *i.e.,* not aligned with what would confer increased fitness to the organism. The exact desired biochemical mode of improvement is often undefined, but rather characterized with carefully designed assays. Generating variants with improved performance relies on computationally predicting assay outcomes, which requires training models on existing assay data using supervised machine learning (6; 7; 8).

Advances of machine learning in the field of structural biology have led to unprecedented accomplishments, like the prediction of structure from sequence (9), sequence from structure (10), the joint generation of both sequence and structure (11) and recently the joint prediction of biomolecular complex structure (12). However, accurate prediction of the effects of mutations on structure and function remains a challenge (13; 14). Pak et al. tried to use AlphaFold to infer the effect of mutations on protein stability and found that it holds little predictive power for this task (15). This result is unsurprising as AlphaFold is trained to predict the structure of a sequence under the implicit assumption that the sequence exists in nature and has a well-formed structure. This assumption is violated when inferring the effect of arbitrary mutations. Although language models are capable of predicting the effect of mutations on native properties (16), their training objective is similarly misaligned with the objective of protein engineering by considering only natural sequences. Another important complicating factor for VEP models is that while a protein may be argued to have one crystal structure (even though this ignores protein dynamics), there is no single unique “stability” or “activity” measure as these depend on precise definitions and assays. As a consequence, large and well-aligned protein function datasets analogous to The Protein Data Bank (PDB) for protein structures are difficult to realize.

In order to improve VEP models, it may help to better understand the aspects that models fail to capture when they make errors. Such insight may enable us to provide our models with the necessary inductive biases and incorporate additional training modalities, such as structural, dynamic or physicochemical features, that better capture these mechanisms, ultimately leading to more efficient protein engineering practices.

As a step in this direction, we seek to identify when VEP models fail to generalize to new, unseen data: what makes the effects of some mutations harder to predict than others? Our analysis focuses on relating the predictability of enzyme variant properties to the structural characteristics of the positions of the mutations. Knowing what determines predictability is useful, because this information can be used to propose variants that models are less likely to make errors on, thereby increasing the detection rate of enzymes with improved properties. Furthermore, it can help improve property prediction models and support benchmarking machine learning models on sequences known to have low predictability, much like how proteins with remote structural homology to training data can be used to assess structure prediction models (17). Lastly, a better understanding of predictability can inform training set design by including larger numbers of variants that are likely to be more challenging — i.e., informative.

To this end, we designed a framework to select positions for mutation with different structural characteristics, and used these to contrast variants with mutations that exhibit those characteristics with variants harboring mutations that do not. In particular, we compared variants with mutations that are 1) buried or exposed, 2) closely or loosely connected with other residues, 3) close to or distant from the active site and 4) part of helices/sheets or part of loops. We hypothesized that mutations that have large effects on the structure and function are more difficult to model and that those mutations are largely defined by their structural characteristics. Buried mutations may cause significant displacement, closely connected mutations are subject to many interactions, mutations close to the active site may perturb the activity to a larger extent and mutations in helices or sheets may disrupt conserved structural patterns.

For each of these four structural characteristics of enzyme mutations, we ask: “Is there a difference in predictability of variants with mutations that have this structural characteristic when compared to variants devoid of those mutations?”. We answered this question by selecting a natural alpha-amylase enzyme (UniProt ID: AMY_BACSU) with potential use for stain removal of starch residues in laundry applications, experimentally creating a large set of variants with multiple mutations and measuring their activity. We repeated this analysis on 12 previously published single mutation enzyme variant datasets to establish whether trends in combinatorial variants, *i.e.,* variants with multiple mutations, and single variant datasets are in agreement or not, and to determine the extent to which predictability is specific to the assay.

## 2 Results

Here, we investigate the influence of different structural properties of mutations on the predictability of their effect, in the context of engineering enzymes for stain removal. We do this by creating a dataset specifically for this purpose, composed of variants of an alphaamylase that all belong exclusively to one structural class. We then test the stain removal activity of these variants and analyse the relation between mutation characteristics and predictability of stain removal activity by machine learning. To our knowledge, no dataset is currently available that allows multiple classes of high-order combinatorial variants — *i.e.,* with more than two mutations — that do not share mutated positions to be partitioned *and* contains a sufficient number of samples for the training of machine learning models.

Next, we set out to assess whether the effects we find occur in isolation or whether it is the introduction of multiple mutations in certain classes that causes differences in performance; and to what extent predictability dependencies are specific to the assay. We therefore repeat our analysis on a site saturation mutagenesis (SSM) dataset of the same enzyme with trisaccharide hydrolysis measurements (18; 19; 20), an SSM dataset of a serine protease of *Cellulomonas bogoriensis* (UniProt ID: A2RQE2_9CELL) with keratin proteolysis measurements (21), and nine publicly available single mutation VEP datasets of different enzymes (S1) from the ProteinGym repository (22). These variants will be referred to as single variants.

### 2.1 An enzyme variant dataset partitioned by structural characteristics of mutations

We consider four types of structural characteristics: buriedness; number of contact residues; distance to the active site; and presence of secondary structure. The aim is to create a dataset that can be split in half based on every characteristic. Henceforth we shall refer to the splits for every structural characteristic as “structural classes”, *e.g.,* the structural class of variants with only buried mutations. The dataset design is visualized in Figure 1(b). In short, we assigned every position binary labels for each structural characteristic that indicate whether it displays that characteristic to a low (0) or high (1) degree. We proceeded by creating sets of positions for every unique combination of structural characteristics, 2^4^ = 16 in total. Variant sequences were then obtained by sampling up to eight mutations per variant from the positions in each set individually, resulting in 16 bins of variants with unique combinations of structural characteristics. These sequences were later cloned and screened, resulting in a dataset of 3,706 variants with associated stain removal activity measurements. Further details on the creation of the dataset are presented in the Methods.

**Figure 1:**
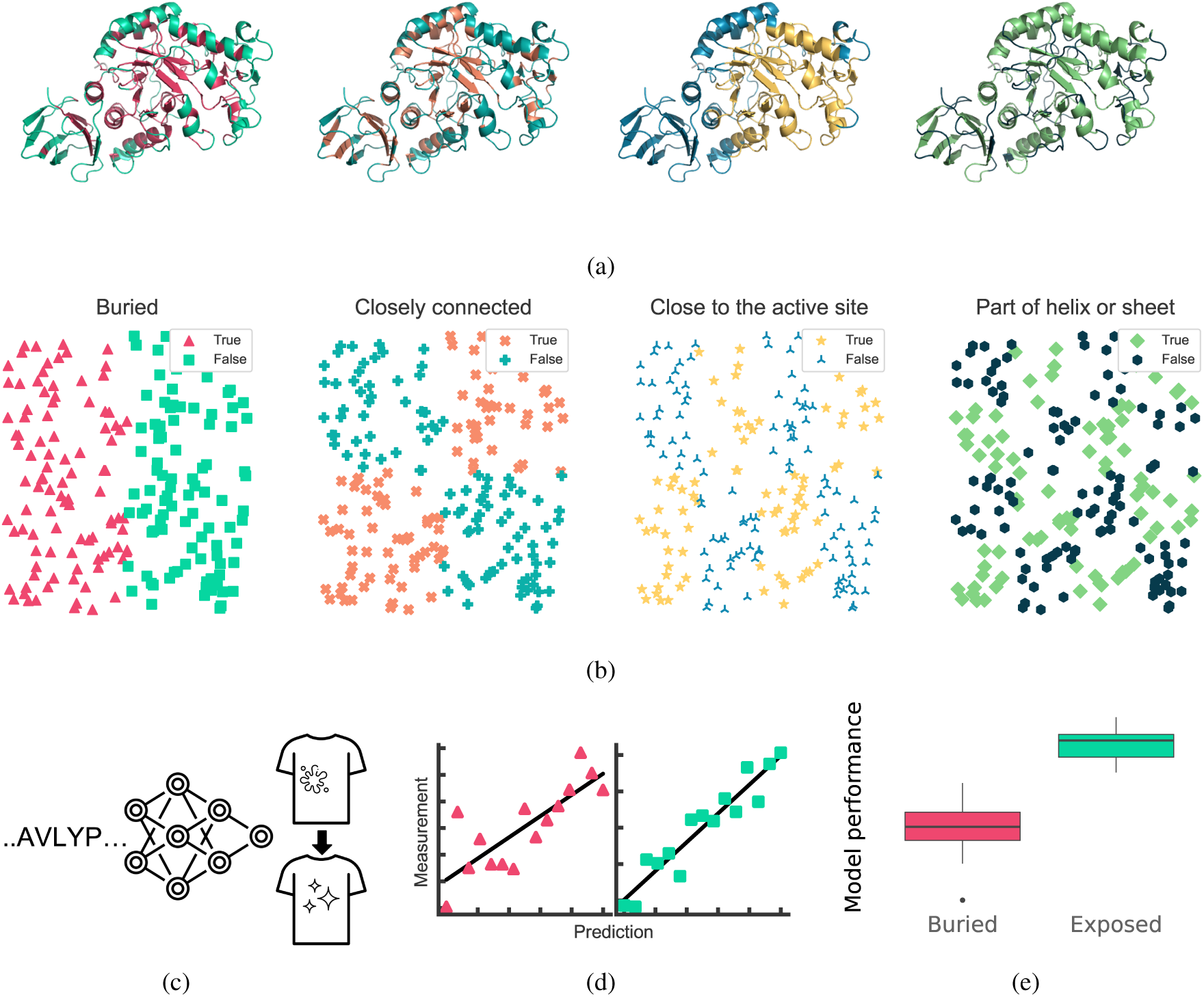
Overview of the approach. (a) We select mutations based on four structural characteristics: buriedness; number of contact residues; distance to the active site; and presence of secondary structure. Every mutation position was then assigned “positive” or “negative” for each characteristic. (b) Illustration of the different splits created by our sequence sampling method. Variant sequences were designed by sampling mutations such that the resulting set of sequences can be divided in two parts of roughly equal size that are positive and negative for each structural characteristic, but also such that there exists a partition of the data for each possible combination of structural classes. These variants are cloned, expressed and screened for stain removal activity. Data shown in the scatter plots are synthetic and axes represent arbitrary sequence projection axes. (c, d) Machine learning models are trained to predict the stain removal activity of the variants based on their sequences. Every structural class of variants is individually trained and evaluated. (e) The performance on held out training data of positive and negative samples is then compared to determine the influence of each characteristic on predictability.

### 2.2 Beneficial and detrimental mutations are found everywhere

We initially set out to determine whether all structural classes contain variants with improved properties. To this end we plotted stain removal activity for all structural classes. The results, shown in Figure 2, show that property distributions are class specific. While activity differences between variants with mutations at helices or sheets and loop regions are not strong, variants with only buried mutations often have minimal activity when compared to variants with exposed mutations, possibly occurring due to conformational rearrangements and loss of stability (23). Despite the differences in activity distributions, all classes contain some variants with higher activity than the reference, which is indicated by the dashed line. This demonstrates that the structural characteristics of mutations are weak predictors of their effects, and that using such characteristics to limit the scope of positions to mutate potentially restricts the identification of variants with improved properties. It is therefore of interest to understand if predictability depends on these characteristics, such that we can focus model improvement on less predictable classes of variants.

**Figure 2:**
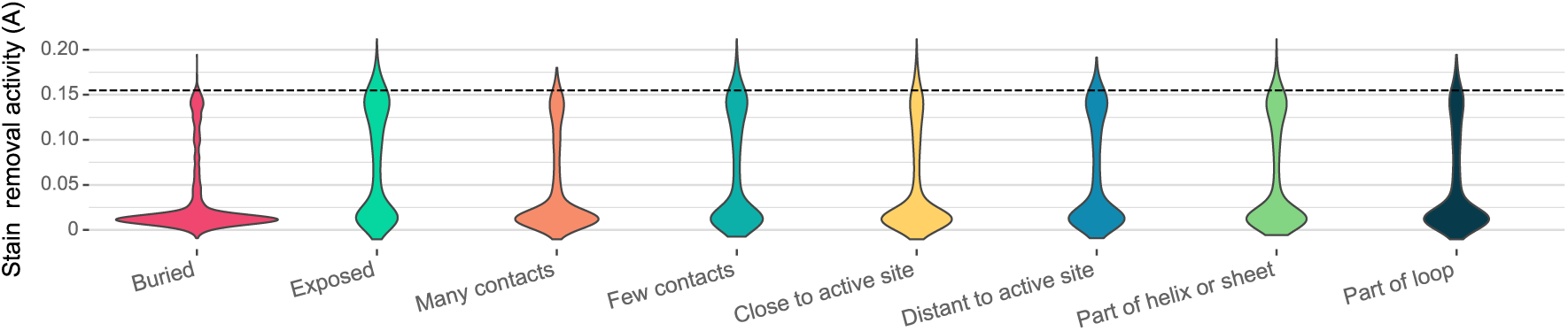
Violin plots of stain removal activity measured by absorbance (A) for every structural characteristic. Stain removal activity of the reference enzyme is indicated by the dashed black line. Every structural split of the data contains improved variants.

### 2.3 Effects of combined mutations with specific characteristics are harder to predict

Next, we examined how these structural characteristics correlate with the predictability of variant effects on activity. Differences in prediction performance for a given split provide insight into the difficulty of modelling the corresponding structural characteristic of mutations. The factorial design of the dataset enabled us to train and evaluate models separately on all structural characteristics, and compare the performance between models trained on both splits individually for a given structural characteristic, *e.g.,* buried and exposed. To minimize the potential for confounding effects, our cloning strategy ensured that all structural classes have a comparable number of samples and a similar distribution of the number of mutations (Table 2) and degree of diversity within each structural class, as measured by edit distance (Supplementary Figure S14).

**Table 1:** Four models are selected with different encoding strategies. Pretraining refers to the data used to train the embedding models. *L*: maximum length of enzyme sequence.

|  | Encoded by | Pretraining | Dimensionality |
| --- | --- | --- | --- |
| **Residue agnostic regressor** | Residue | None | $L$ |
| **Partial least squares** | Residue | None | $L \times 20$ |
| **Augmented Potts model** | Residue | MSA | $L \times 20 + 1$ |
| **RITA regressor** | Sequence | UniRef-100 + MGnify<br>+ Metaclust | 4096 |

**Table 2:** Number of variants in the combinatorial dataset per structural class in total and per number of mutations.

| Structural characteristic | Number of Variants |  |  |  |  |  |  |  |  |
| --- | --- | --- | --- | --- | --- | --- | --- | --- | --- |
|  | Total | 1 Mut. | 2 Mut. | 3 Mut. | 4 Mut. | 5 Mut. | 6 Mut. | 7 Mut. | 8 Mut. |
| Exposed | 1,823 | 254 | 519 | 504 | 320 | 168 | 46 | 10 | 2 |
| Buried | 1,883 | 235 | 528 | 561 | 354 | 142 | 47 | 16 | 0 |
| Few contacts | 1,905 | 232 | 490 | 532 | 374 | 202 | 61 | 12 | 2 |
| Many contacts | 1,801 | 257 | 557 | 533 | 300 | 108 | 32 | 14 | 0 |
| Distant to active site | 1,924 | 255 | 537 | 574 | 343 | 157 | 45 | 12 | 1 |
| Close to active site | 1,782 | 234 | 510 | 491 | 331 | 153 | 48 | 14 | 1 |
| Part of loop | 1,747 | 248 | 506 | 501 | 313 | 125 | 42 | 11 | 1 |
| Part of helix or sheet | 1,959 | 241 | 541 | 564 | 361 | 185 | 51 | 15 | 1 |

Figure 3 shows large differences in median performance between variants positive and negative for the characteristics buriedness, number of contact residues and distance to the active site. Combinatorial variants in our dataset appear more difficult to model when their mutated sites are buried, close to the active site or in contact with many residues. We observe slightly better predictions for mutations occurring at helices or sheets versus loop regions.

The results shown in Figure 3 are almost identical across all models, which indicates that predictability differences are nearly independent of model type for the combinatorial dataset. Although evolutionary models and language models produce rich sequence features and model mutations in the context of all other residues, the mechanism by which mutations affect stain removal activity in less predictable variants is apparently not well accounted for by these models.

**Figure 3:**
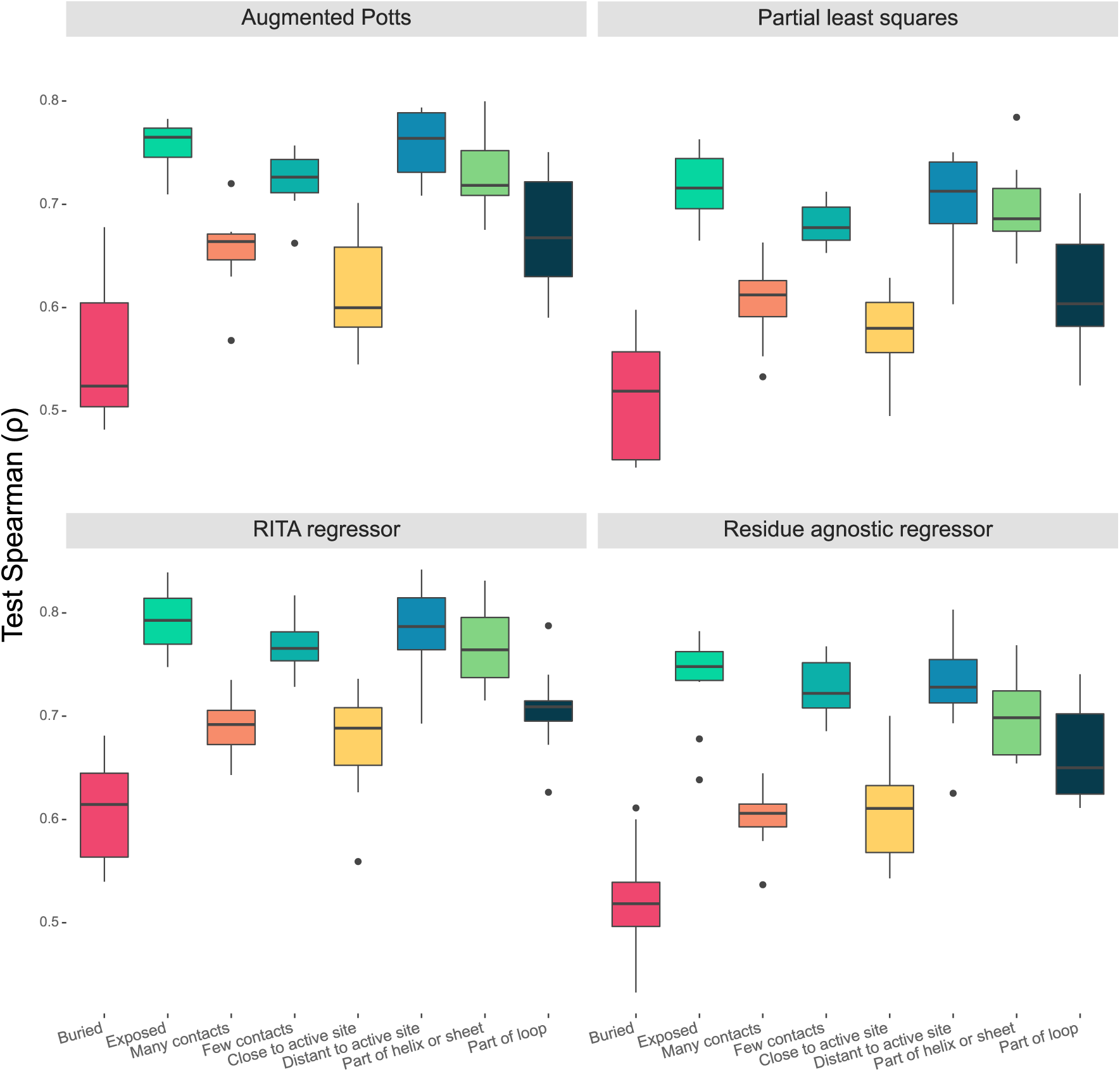
Test Spearman correlation score distributions of all four models on 10 folds of the combinatorial training data. The models exhibit significantly worse prediction performance on combinatorial variants with mutations at sites that are buried, close to the active site, and in contact with many residues.

### 2.4 Single mutation variant effect predictability depends on the same structural characteristics

The findings from the combinatorial variant analysis led us to explore whether similar patterns exist in single mutation variants. We used a dataset of the same enzyme, as well as a dataset of a *Cellulomonas bogoriensis* serine protease, both with *in vitro* measurements, and a collection of public datasets that measure enzyme activity both *in vivo* and *in vitro*. Interestingly, in most of the datasets, we identify the same structural dependencies we observe in the combinatorial datasets, but with different effectdirections, as shown in the boxplots in Figure S1-S10. More importantly, despite choosing four models with different feature extraction methods, the parallel coordinates plot in Figure 4 shows that predictability differences are highly similar across models, demonstrating that these differences are primarily driven by the assay, and less so by the models.

**Figure 4:**
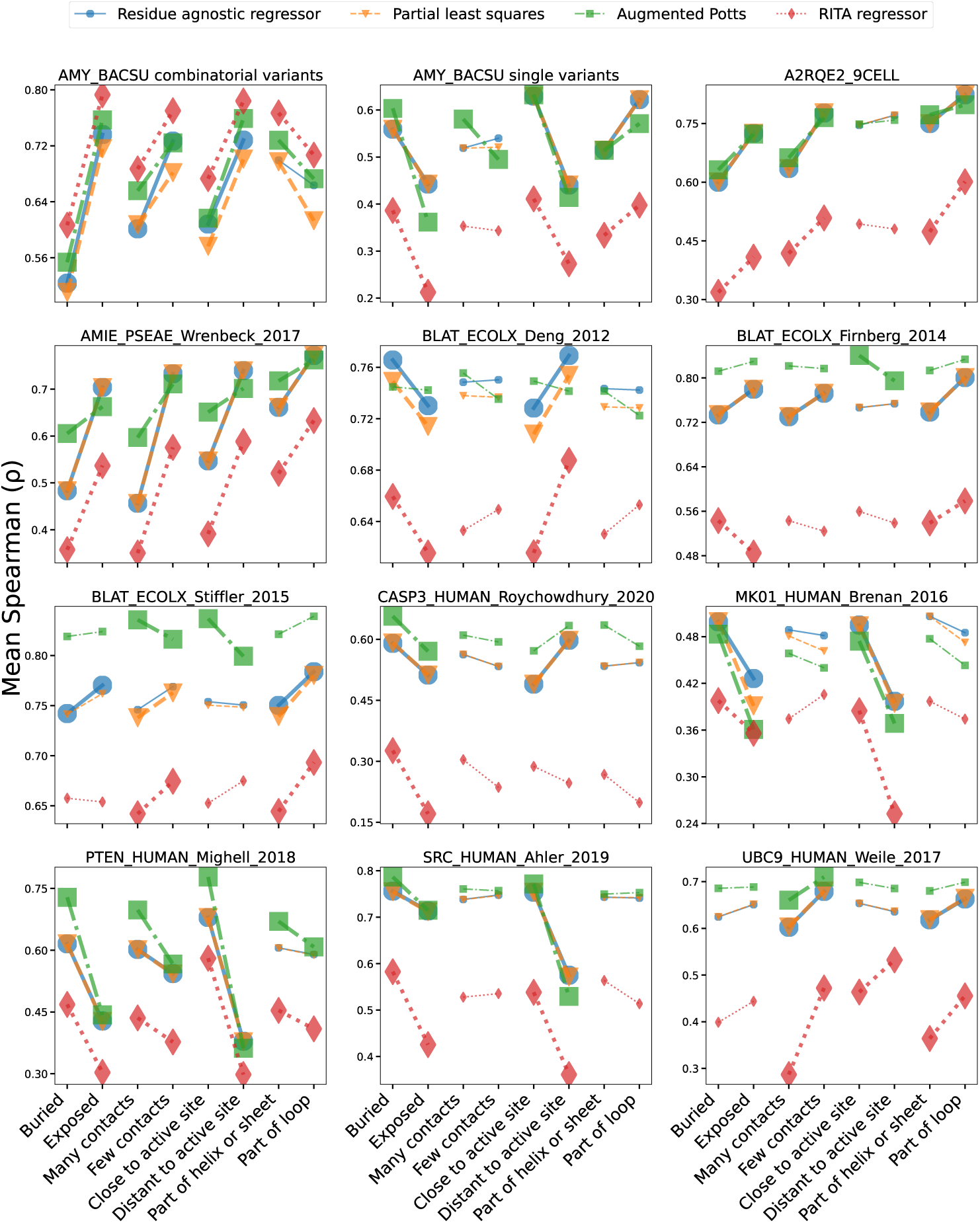
Parallel coordinates plot of average Spearman scores, illustrating similarity in variant predictability of all structural classes between all four models. For a given pair of opposing structural classes, *e.g.*, buried and exposed, the slope similarity of any two models’ line segments represents their similarity in predictability differences for those structural classes. Line segments with increased width denote predictability differences that are statistically significant, as determined by an independent samples t-test. Studies corresponding to the public datasets are organized in Table S1.

## 3 Discussion

In this study, we have shown that the difficulty of variant effect prediction in enzymes correlates systematically with structural features. As a specific example, we created a balanced dataset for an alpha-amylase and observed that variants with mutations at sites that are buried, in close contact with other residues and close to the active site are harder to predict than their counterparts for baseline models, as well as more complex pretrained models. These dependencies also presented themselves in single mutation variants, but with directions that were specific to the assay and mostly unaffected by the type of model.

These findings are somewhat surprising, since both the augmented Potts model and the RITA regressor rely on different features than our baseline models. This can be deduced from their performance on variants with mutations at positions not mutated in the training data (22), and we corroborate this in S12. Although the augmented Potts model sometimes performs better on classes with low predictability, models generally make similar mistakes, suggesting that different modalities of protein variants and model architectures with new inductive biases will play important roles in further improvement of variant effect prediction.

Further insights could be obtained by evaluating new prediction strategies on all structural classes of datasets that display strong predictability differences. We assume that all classes of variants are affected by the same noise distributions, caused by stochastic and systematic error. A perfect model should therefore perform equally well on positive and negative classes of variants, which can be evaluated by explicitly using them as individual benchmarks.

### 3.1 Limitations of existing protein variant datasets

To date, there are many variant datasets for various proteins and accompanying properties, that resulted from different methods of sampling the mutational landscape (22). The majority are comprised of single mutation variants, although some datasets are combinatorial, *i.e.,* contain variant sequences with more than one mutation. Some protein variant datasets are limited to a small subset of residue positions (24), while others have no constraints on the positions from which mutations are sampled (25).

An inherent limitation of combinatorial protein variant datasets to address the research question we focused on, is that stratification by mutated positions is hindered by variants with overlapping mutated positions. For instance, when comparing variants with mutations exclusively in the core of the protein with those with mutations only on the surface, all variants that contain mutations both in the core and on the surface should be disregarded.

An example of a dataset with non-overlapping mutated positions was recently published by Tsuboyama *et al.* (26). This dataset measured the folding stability of both single and double mutants of 331 natural and 148 *de novo* designed proteins. While the dataset presented here is far smaller, it is of an enzyme of significant size (425 aa) instead of single domains, with higher-order combinatorial variants and mutations covering nearly all residues as opposed to small subsets – all features often encountered in protein engineering.

Our dataset is unique in its potential for stratification by mutation characteristic, while also being an exemplar of a typical protein engineering dataset encountered in industrial scenarios. We hope that the dataset, as well as the design strategy, will prove useful for advancing machine learning guided protein engineering.

### 3.2 Challenges and prospects for future research

One of the central challenges in protein engineering is predicting the fitness of variants that lie beyond the training data domain. To design proteins with improved fitness, it is essential to predict the effect of new mutations as well as properties that exceed the domain of the training data. Fannjiang and Listgarten (27) make the case that both training set design and providing models with domain-specific inductive biases hold the most potential in achieving good extrapolation to novel mutations and improved properties. This assertion is supported by our analysis, which establishes that predictability dependencies on structure depend less on model choice than on the training data at hand. This suggests that there are characteristics of the mechanisms that relate sequence to property that are consistently not captured by the models included in our analysis. One way of addressing this issue is to include stronger biological priors into VEP models. Proteins have been successfully engineered with mechanistic models (28; 29), which rely on physical information that is not explicitly represented by sequence models and is free of evolutionary bias. Adding explicit physical features to models, or imposing constraints directly into the model architecture with so-called physics informed neural networks, could be an approach to improve performance on less predictable classes of variants.

Other than addressing model capacity to extrapolate, there are also challenges inherent to the training data. The design space of single mutation variants is relatively small, and can in many cases be exhaustively experimentally characterized for a given property of interest. The combinatorial design space on the other hand is many orders of magnitude larger, and mutations often do not modulate the property in isolation, but rather display interaction effects resulting in a phenomenon that is referred to as epistasis. How epistasis influences predictability is not clear and establishing this is difficult, if only because epistasis of protein mutations is not clearly defined. The widely used definition “non-linear interaction of mutations” oversimplifies the mechanisms that couple sequence to property, since most properties are non-linear in nature to begin with, *e.g.,* due to the existence of upper and lower bounds and diminishing returns in substrate turnover caused by rate limiting factors such as substrate concentration and diffusion. Methods exist that aim to quantify position-specific epistatic terms (30), and an interesting follow-up would be to investigate whether these terms can be coupled to predictability by machine learning models, and how these terms are distributed over structural classes of enzyme variants.

Another obstacle relating to the training data is the domain shift induced by generation of successive screening data. Sequences of successive screens are often not sampled from the same distribution as the training data, as most protein variant datasets are generated with the purpose of property optimization.

Finally, there is still room for improvement in performance evaluation. New methods in machine learning guided protein engineering are often evaluated by their rank correlation on randomly held out data, which does not satisfactorily address the capacity to extrapolate. Dallago et al. have proposed strategies of evaluating extrapolation by creating data splitting strategies based on the dependent variable and number of mutations (31), and recently, Notin et al. have proposed the “modulo” and “ contiguous” data splits (22), which assure mutually exclusive mutated positions in the splits. Such splitting strategies have the potential to advance the field of VEP, but are not always applicable to combinatorial datasets. Our work demonstrates how dataset design plays an important role in illuminating data characteristics that cause modelling challenges, and provides data for targeted model improvement.

## 4 Methods

### 4.1 Data

#### 4.1.1 Sampling of combinatorial variant sequences

We created a dataset of variants with multiple mutations that can be stratified into two sets of roughly the same size for four structural characteristics. The number of samples per structural characteristic of mutations is described in Table 2.

##### Calculation of structural characteristics for each position

We started by computing all structural characteristics of the alpha-amylase using PDB structure 1UA7 (32), the latest out of the two available structures for the amylase we measured, for every residue on a continuous scale. Buriedness was quantified as the shortest distance of the distances of all atoms in the residue to the convex hull of the enzyme, which was computed using the ConvexHull class of the Spatial module of scipy version 1.9.2. The degree to which residues are connected was computed by counting the number of *C_α_* atoms within a given radius from the *C_α_* atom of the considered residue. This radius was set to the average distance of neighboring *C_α_*atoms plus two corresponding standard deviations, which in the case of this enzyme resulted in 7.3 Ångström. The distance to the active site was found by computing the distances of a residue *C_α_* atom to the *C_α_* atoms of the two residues at the active site (D176, E208), and then taking the shortest of these distances. Lastly, secondary structure was assessed with the ProDy python package version 2.1.2. (33). Structural characteristics of the enzymes from the remaining datasets were assigned as described above, using PDB structures located at https://github.com/florisvdf/mutation-predictability.

##### Assignment of binary structural characteristics to positions

To turn these continuous structural characteristics into binary values, we computed the median of the respective characteristics and then assigned them a binary label using this median as a threshold. In the case of buriedness, all positions with a value higher than the median were assigned a positive label and negative otherwise. Positions were assigned a positive label for “many contacts” if they had more contacts than the median number. Positions closer to the active site than the median distance were assigned a positive label for being “close to the active site” and negative otherwise. Finally, secondary structure labels were assigned based on the secondary structure assessment as described in the previous paragraph. While buriedness and number of contact residues may seem like similar concepts, in Figure S11 we demonstrate that the marginal probabilities of residues being exposed and closely connected and residues being buried and loosely connected are substantial, which drove us to include both characteristics in our analysis separately.

##### Grouping of positions

To ensure that we can split our data for every structural characteristic, we opted for a factorial design strategy: for every unique combination of structural characteristic labels, we grouped the positions into sets associated with a bin. Every bin contains variants with mutations that occur only at positions of the associated set. This results in a total of 4^2^ = 16 bins. Distributions of the structural characteristic values per bin are presented in Supplementary Figure S13.

##### Sampling of variant sequences

The final step in creating a dataset of variant sequences that can be split for every structural characteristic was to sample variants from each bin. To obtain a near equal number of variants for all negative and positive splits, we sampled equal numbers of variants from all 16 bins. For a given bin, variants were generated by uniformly sampling from each set of positions and mutating residues randomly. We sampled residues of 75% of the variants from protein language model (PLM) likelihood scores of ESM1v (esm1v_t33_650m_ur90s_1) in an effort to optimize the yield of the cloning pipeline, as such scores have been shown to correlate with expression (16). The remaining 25% of variants were sampled uniformly over all possible alternative residues, ensuring that some fraction of the variants are not constrained to diversity that is biased towards higher PLM likelihoods. This yielded 21,436 candidate alpha-amylase variants, of which 3,706 were successfully synthesized, cloned and transformed (see below). Variant sequences of the resulting transformants were later tested for stain removal activity.

#### 4.1.2 Cloning

Expression cassettes encoding the enzyme variants were constructed by ordering DNA oligos representing all 21,436 candidates from Integrated DNA Technologies and assembled by PCR. The resulting fragments were fused to a signal sequence, ATGAAACAACAAAAACGGCTTTACGCCCGATTGCTGACGCTGTTATTTGCGCT-CATCTTCTTGCTGCCTCATTCTGCAGCTTCAGCAGAAACGGCGAACAAATCGAATGAG, to drive secretion of the amylase, and flanking sequences to allow for integration into the genome of *Bacilus subtilis* upon transformation. The expression vector of obtained transformants was PCR amplified with barcoded primers and resulting amplicons were sequenced using Oxford Nanopore Technologies for validation. 3,706 variants that matched the intended designs were picked and grown in 96-well format. Culture supernatants containing the secreted amylase variants were used for downstream assays without further purification.

#### 4.1.3 Stain removal screening

Enzyme solutions were screened for stain removal in triplicate on 96-well plates that were prepared with swatches of fabric with starch-based stains (Rice Starch, CS28 https://www.cftbv.nl/) applied (34). Non-normalized stain removal activity of all alpha-amylase variants was measured as released soil through absorbance measurements of the supernatant of the incubated swatch-enzyme wells. Final non-normalized activity data were obtained by subtracting a blank absorbance measurement from the raw measurements and taking the average of the triplicate measurements, resulting in values between −1.03 × 10*^−^*^2^ and 2.12 × 10*^−^*^1^. Concentration normalized stain activity data were also obtained, but were not included in this analysis due to technical problems with determining concentration. We pose that non-normalized stain activity is a valid property to evaluate VEP models, since its constituent properties, expression and reaction rate, are both functions of the sequence.

### 4.2 Single mutation variant datasets

The single variant dataset presented here is an SSM dataset. Out of all 8,075 possible variants, 7,467 were successfully cloned and screened for their activity on glucose polymers with a degree of polymerization of 3. These activity values were subsequently normalized by the measured activity of the reference (unmutated) enzyme and used for fitting the regression models. The dataset contains variants with single mutations at each position, with the exception of position 4, 233, 399 and 417 (18; 19; 20).

We selected nine additional SSM datasets of enzymes from the ProteinGym repository that measure activity. We constrained our selection to datasets that contain at least 1,000 single mutants, contain sequences no longer than 600 residues and have a corresponding reference structure that covers the mutated region.

### 4.3 Model definitions

We selected four linear regression models, each trained on embeddings ranging in complexity. To allow for higher model complexity, non-linear relations between in- and outputs and utilization of unlabeled sequence data, we used two types of pretrained models to embed our training data for downstream regression of our linear models - the RITA regressor and the augmented Potts model (35). These models leverage rich representations of our sequences and model interactions of mutated residues in our sequences:

- The augmented Potts model is a ridge regression model trained on a concatenated vector of one hot encoded variant sequences and the sequence energy obtained from a Potts model (36). Potts models learn to reconstruct the sequence distribution of a family of sequences, which in the case of protein sequences are generally organised in a multiple sequence alignment (MSA). This sequence distribution is reconstructed by learning a set of field parameters *h_i_* that represent the propensity of every possible amino acid occurring at every sequence position *i*, and coupling parameters *J_i,j_* that represent the propensity of every possible pair of amino acids occurring at every pair of sequence positions (*i, j*) (37). Sequence energies are computed by summing the field parameters *h_i_* of each residue *σ_i_* in the sequence *σ*, and the coupling parameters *J_i,j_*of each combination of residues (*σ_i_, σ_j_*), where *i* and *j* denote positions in the sequence:

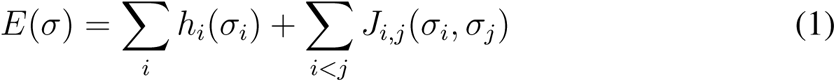

We used MSAs that correspond to the reference sequence from the ProteinGym repository to train our Potts models, and generated MSAs for the alpha-amylase and the *Cellulomonas bogoriensis* serine protease using the same protocol, as described by Notin et al. (38).
- The RITA regressor is a ridge regression model trained on embeddings from an autoregressive language model. Autoregressive language models are commonly used to generate embeddings of input sequences by averaging the hidden states of the sequence over the sequence dimension. This method is effective because each hidden state is conditioned on the previous hidden state, allowing for the modeling of interactions between all tokens in the sequence, and thus introducing sensitivity to interactions between mutations. Note that the hidden state is aggregated over the entire sequence. Features thus no longer correspond to individual residues, but to properties of the entire sequence. The regression model is trained to capture the effect of learned sequence features on the mutation effect, rather than simply additive contributions from individual residues.

These two more complex embedding strategies are compared to two baseline approaches that are only trained on one-hot encodings of the variant sequences. The difference between the two baseline models is that the Partial Least Squares model is trained on one hot encodings of the entire sequences, whereas the Residue Agnostic Regressor is trained on an indicator vector of length *L*, where *L* is equal to the sequence length and the indicator function 1(*σ_i_*) evaluates to 1 when the residue at position *i* in the sequence *σ* is mutated, and 0 otherwise:

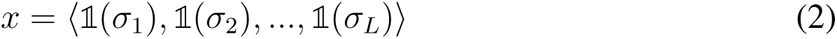

Thus, any two sequences in a protein variant dataset with mutations at the same positions have identical encodings. Comparison to these baselines provides information on the effect of additional signal obtained from residue type on modelling difficulty.

The Residue Agnostic Regressor, augmented Potts model and RITA regressor were all implemented using the Ridge class from the linear_model module of scikit-learn version 1.1.2 (39). The regularization parameter, *α*, was chosen by performing a search over each entire dataset, evaluating *α* values of 0.001, 0.01, 0.1, 0.9, and subsequently choosing a single *α* value that resulted in the best performance across datasets (see Section 4.4), thereby evaluating the models under a best case scenario in our analysis (Supplementary Figures S15, S16). The *α* value had a negligible effect on the augmented Potts model performance, but yielded the best performance for the RITA regressor when set to 0.1, which we used for all three models.

The Partial Least Squares model was also implemented in scikit-learn. We used the PLSRegression class of the cross_decomposition module and set the n_components parameter to 20. The augmented Potts model embeds sequences using GREMLIN_CPP v1.0 with default parameters, except for the gap_cutoff parameter. The Potts model ignores residues in the sequence that have a gap ratio that exceeds the gap_cutoff parameter. We set this parameter to 1 to ensure that emission parameters were computed for every residue in the sequence. The RITA regressor embeds sequences using RITA_xl from the huggingface repository lightonai/RITA_xl (40). RITA_xl produces embeddings with a hidden state dimensionality of 2048 for each sequence direction. We obtained the final embeddings by concatenating the embeddings for both directions, resulting in an embedding dimensionality of 4096.

### 4.4 Model evaluation

For combinatorial variants, we obtained model performance estimates by training on both the negative and positive variant classes separately for all four mutation characteristics, using 10-fold cross validation and taking the mean Spearman correlation coefficients between predicted and measured stain removal activity over all the test-folds as a performance measure.

For the single mutation variants, we made one adjustment to the 10-fold cross validation procedure: instead of randomly sampling test-folds, we sampled folds by taking random fractions of variants with mutations at each site individually, ensuring that no sites are mutated in the test folds that are not also mutated in the corresponding training folds. This method of cross validation ensures that the Residue Agnostic Regressor and Partial Least Squares make informed predictions, as evaluating them on variants with mutations at novel positions will yield predictions that are independent from the inputs. The resulting folds will be similar to random folds, with the only difference being a small fraction of variants with mutations at novel positions not occurring in the test folds, which we assume to have a negligible effect on predictability patterns.

## Acknowledgements

We thank Viktor Alekseyev for reviewing the manuscript and providing important feedback. We would also like to express our gratitude to the members of the Bioinformatics Group at Wageningen University for discussion and input.

## Author contributions

F.J.v.d.F. and H.R. conceived the research project. H.R. conceived, implemented and carried out the stratified dataset design. F.J.v.d.F. implemented and trained the machine learning models, carried out the comparative analysis, created figures, wrote the manuscript and software. A.D.J.v.D, D.d.R. and H.R. supervised the research project and contributed to writing the manuscript. D.E. contributed the data on single mutants and provided supervision. H.M., S.v.S.T, and R.O. created the mutant library. F.J.v.d.F, S.P., L.D., F.G. and D.S. carried out the stain removal activity screens. L.L. performed enzyme concentration quantification by HPLC.

## Competing interests

The authors declare no competing interests.

## Data availability

All data used to generate the results presented in this work is made available at https://github.com/florisvdf/mutation-predictability.

## Code availability

All python code used to produce the results presented in this work is publicly available at https://github.com/florisvdf/mutation-predictability.

## A Supplementary Materials

**Figure S1:**
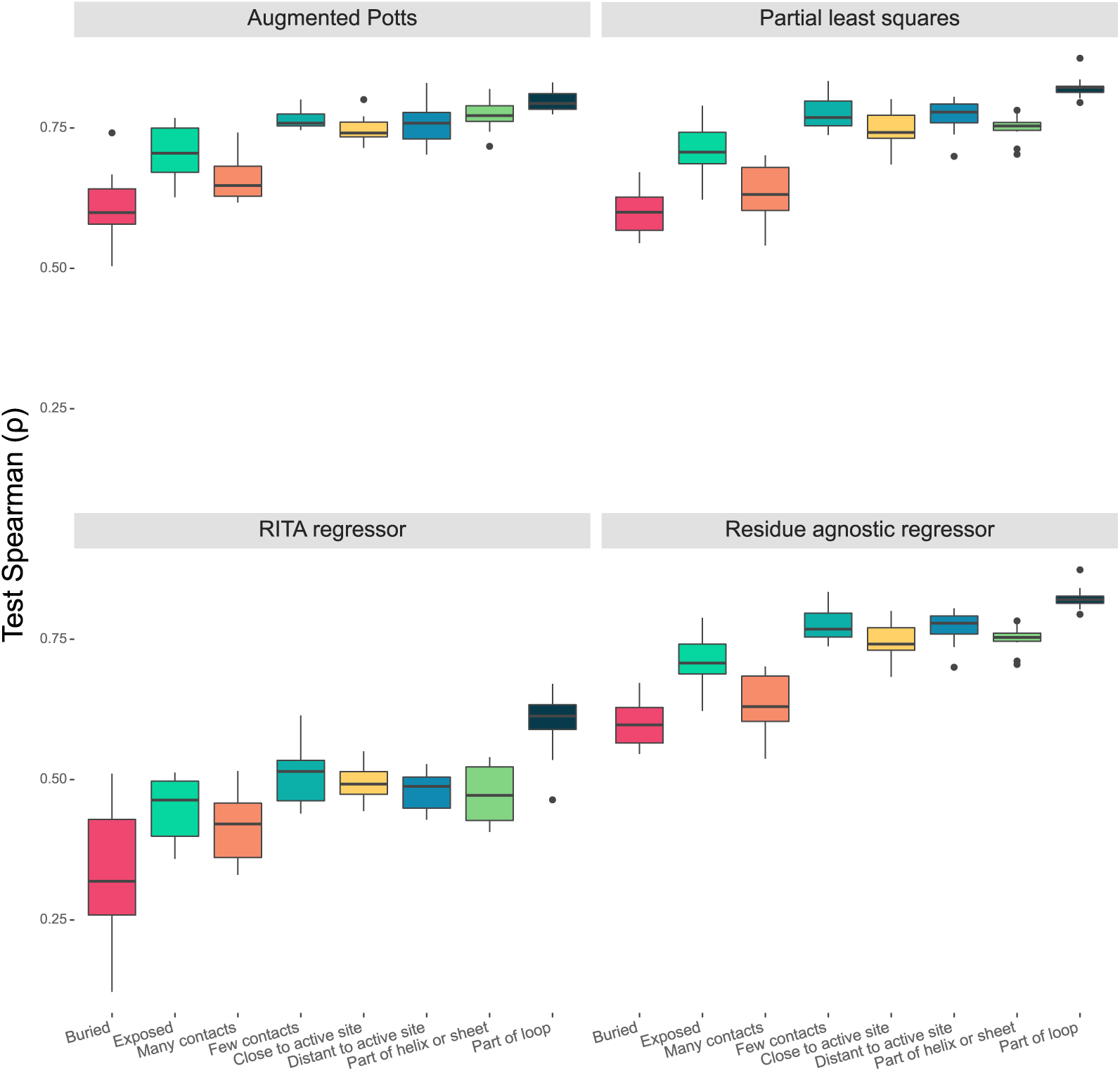
Test Spearman correlation score distributions of the all four models on 10 folds of the A2RQE2_9CEL dataset.

**Figure S2:**
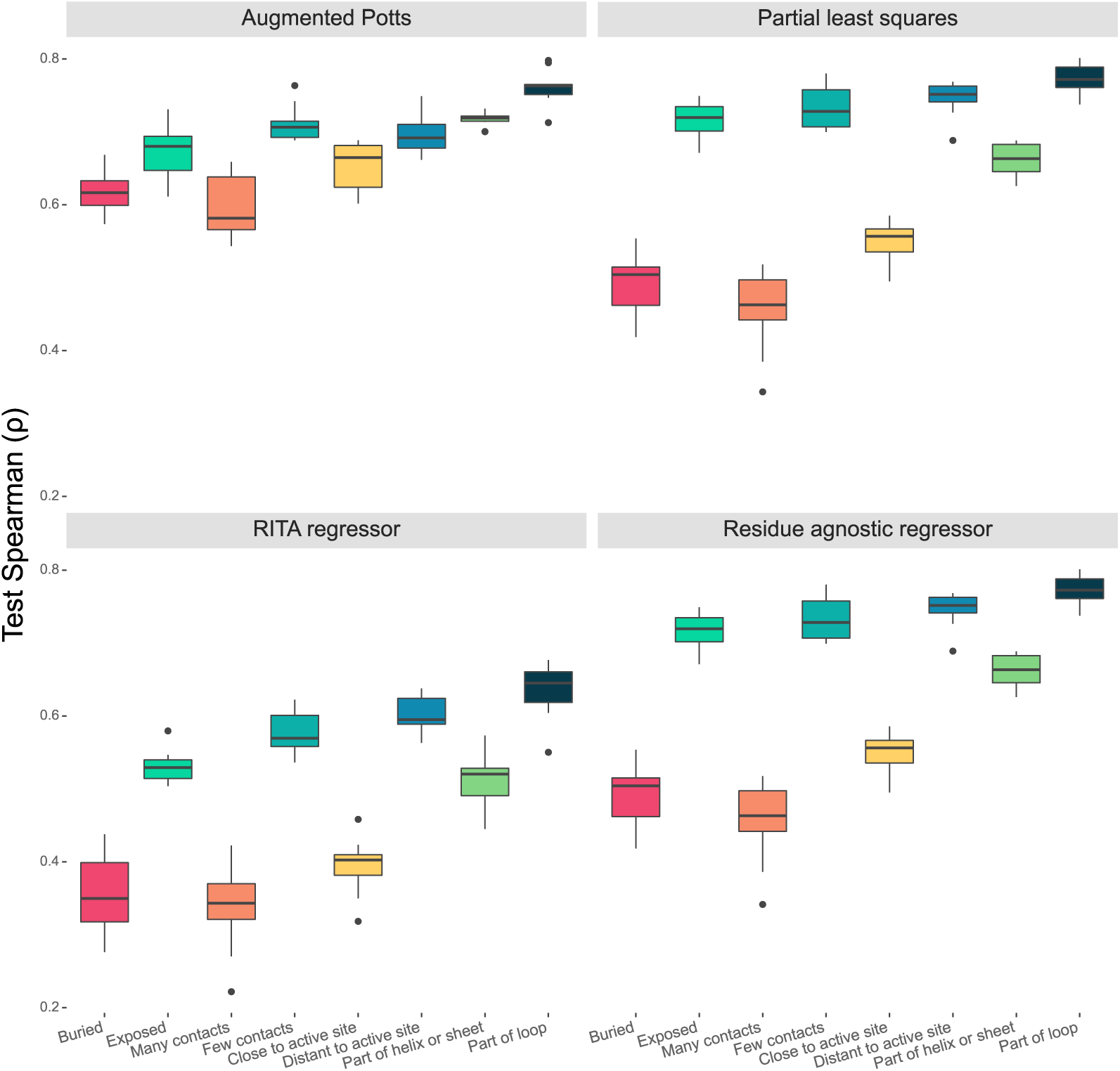
Test Spearman correlation score distributions of all models on 10 folds of the AMIE_PSEAE_Wrenbeck_2017 dataset.

**Figure S3:**
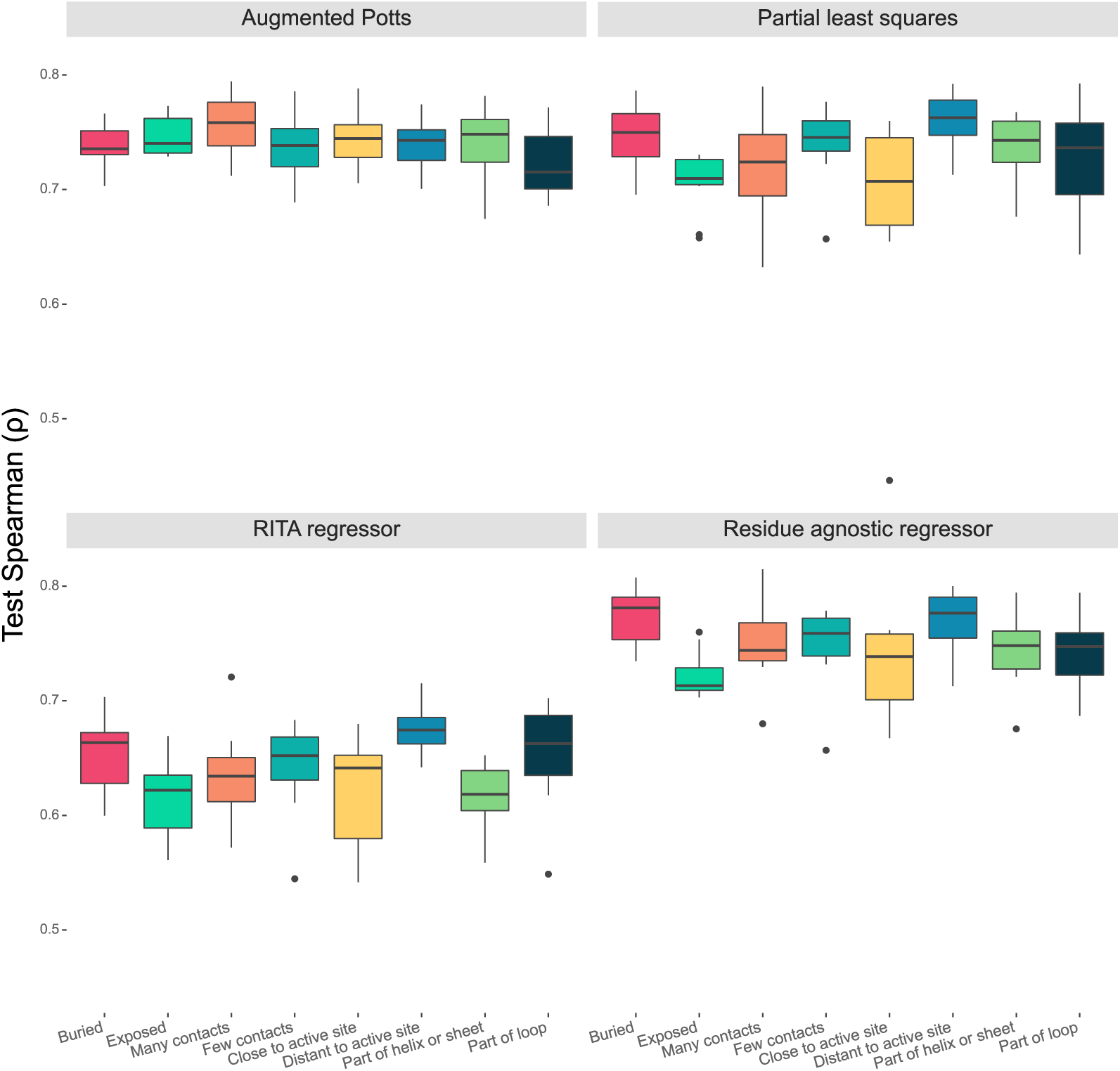
Test Spearman correlation score distributions of all models on 10 folds of the BLAT_ECOLX_Deng_2012 dataset.

**Figure S4:**
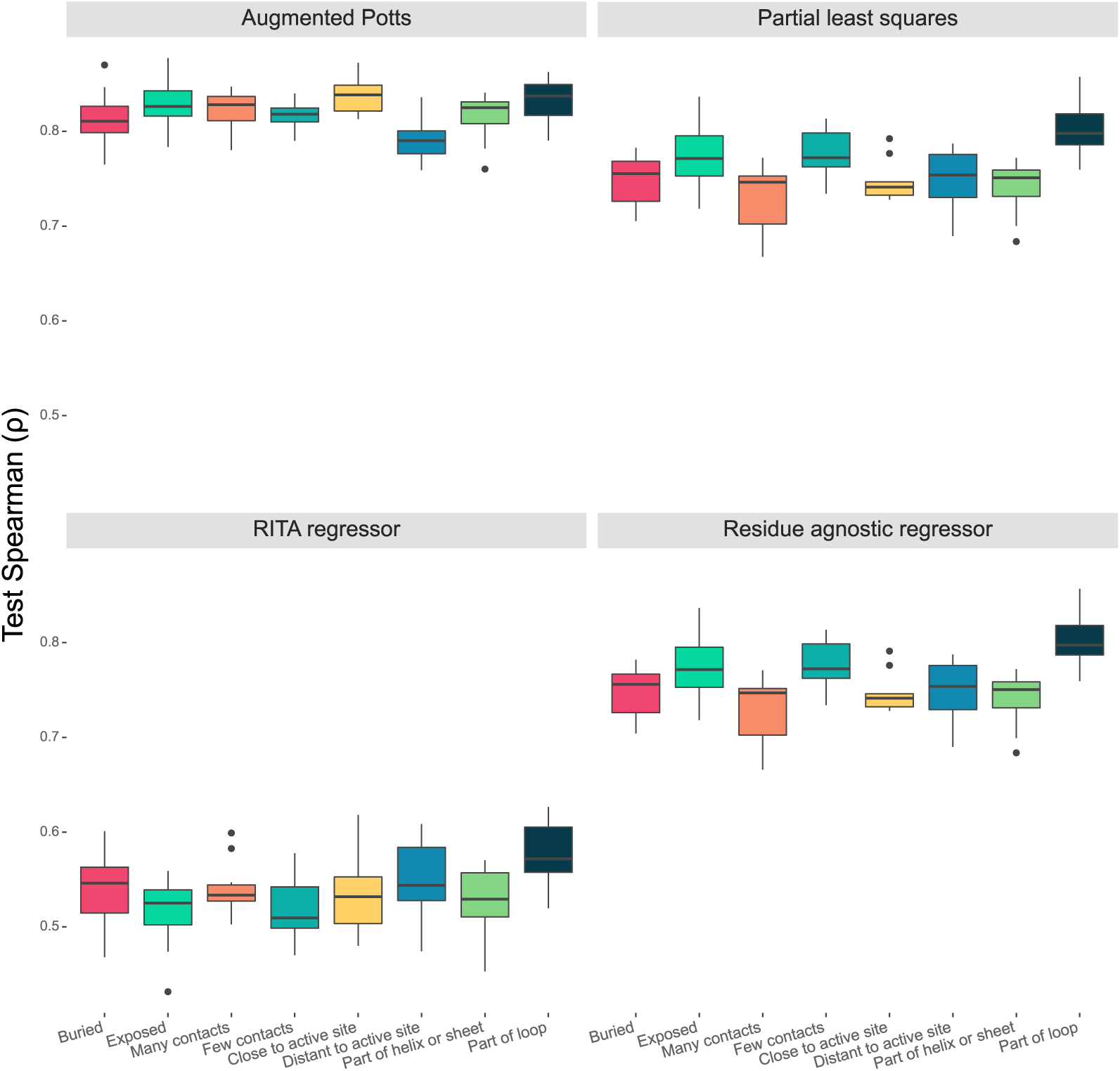
Test Spearman correlation score distributions of all models on 10 folds of the BLAT_ECOLX_Firnberg_2014 dataset.

**Figure S5:**
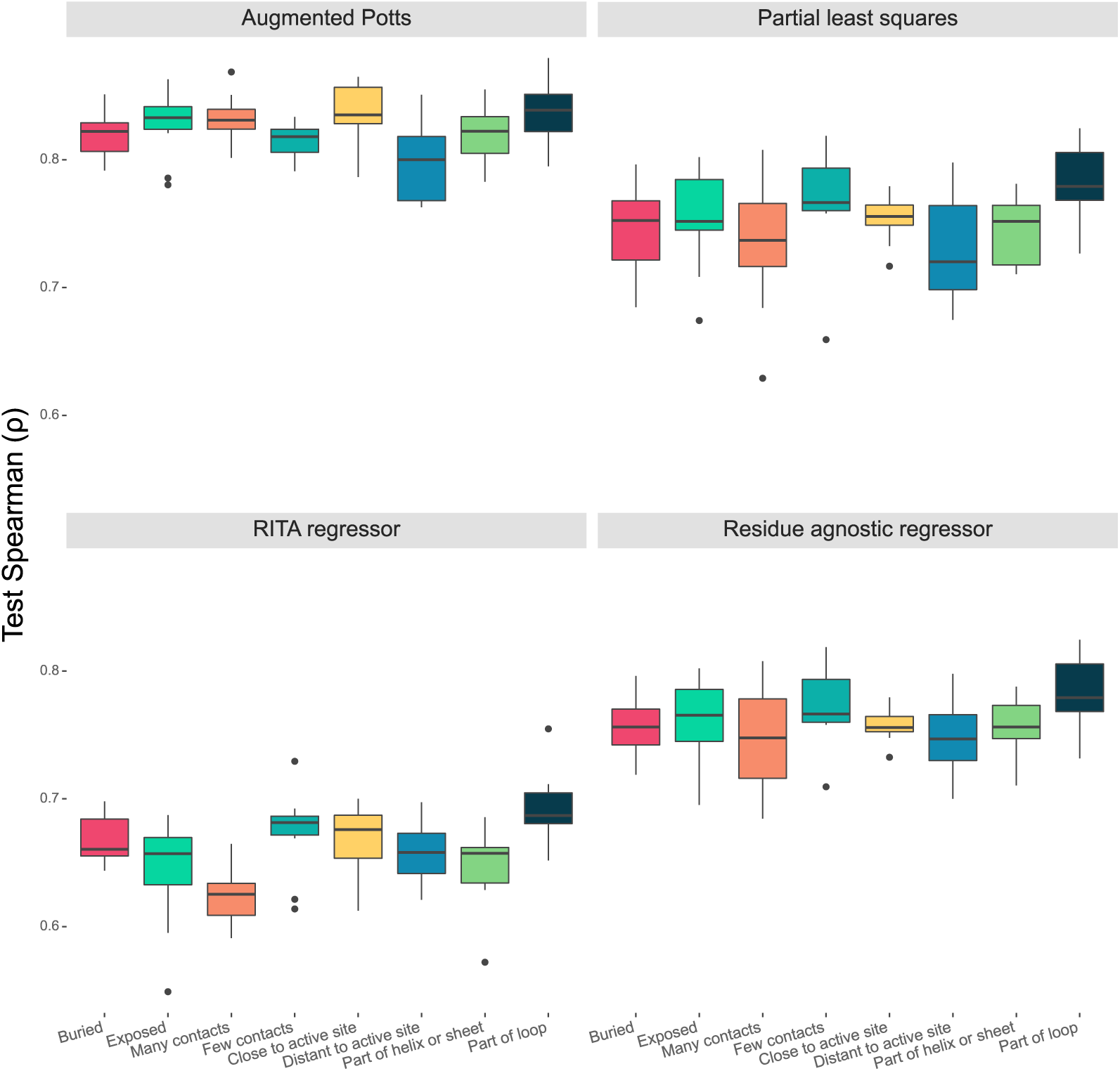
Test Spearman correlation score distributions of all models on 10 folds of the BLAT_ECOLX_Stiffler_2015 dataset.

**Figure S6:**
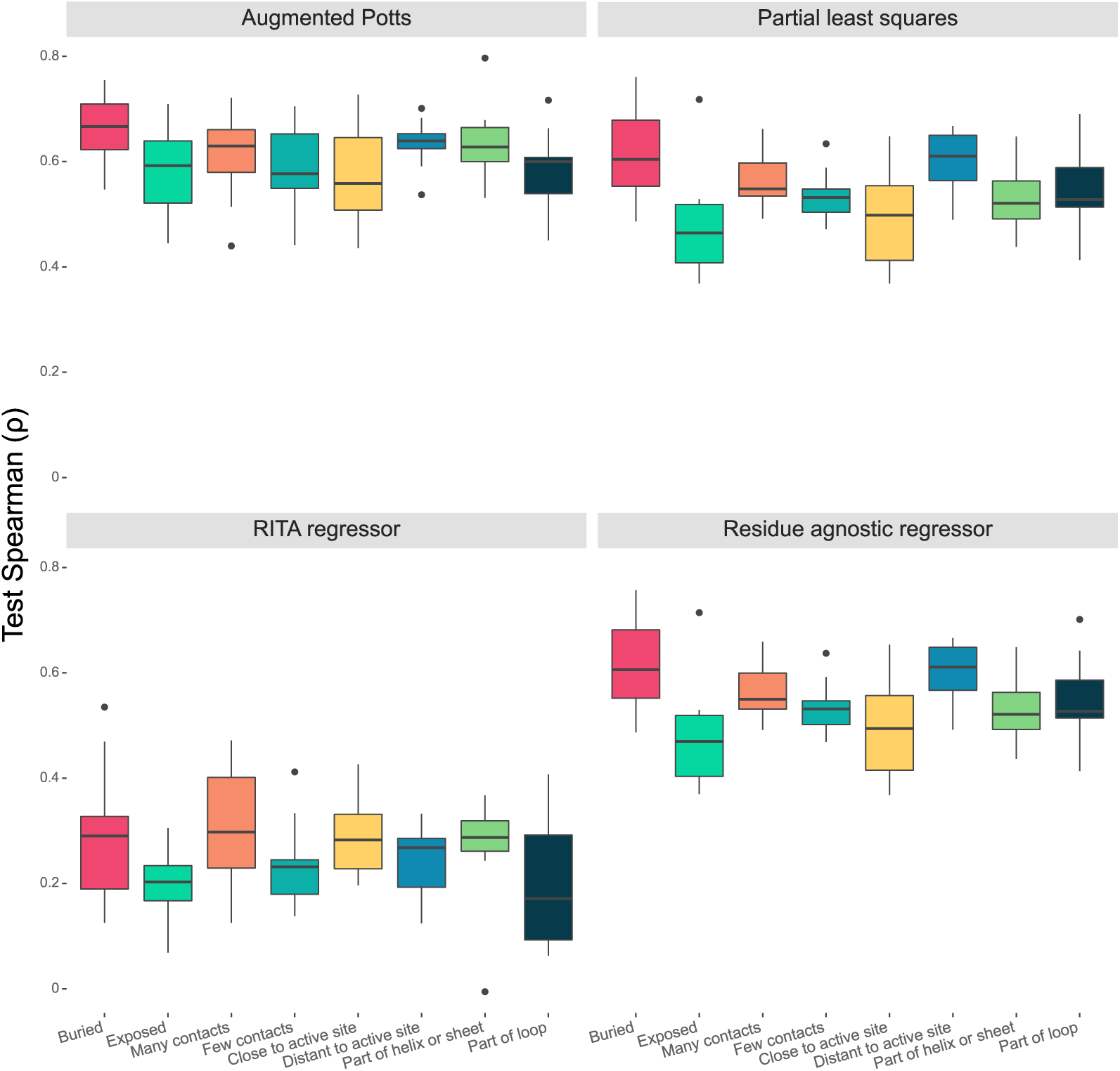
Test Spearman correlation score distributions of all models on 10 folds of the CASP3_HUMAN_Roychowdhury_2020 dataset.

**Figure S7:**
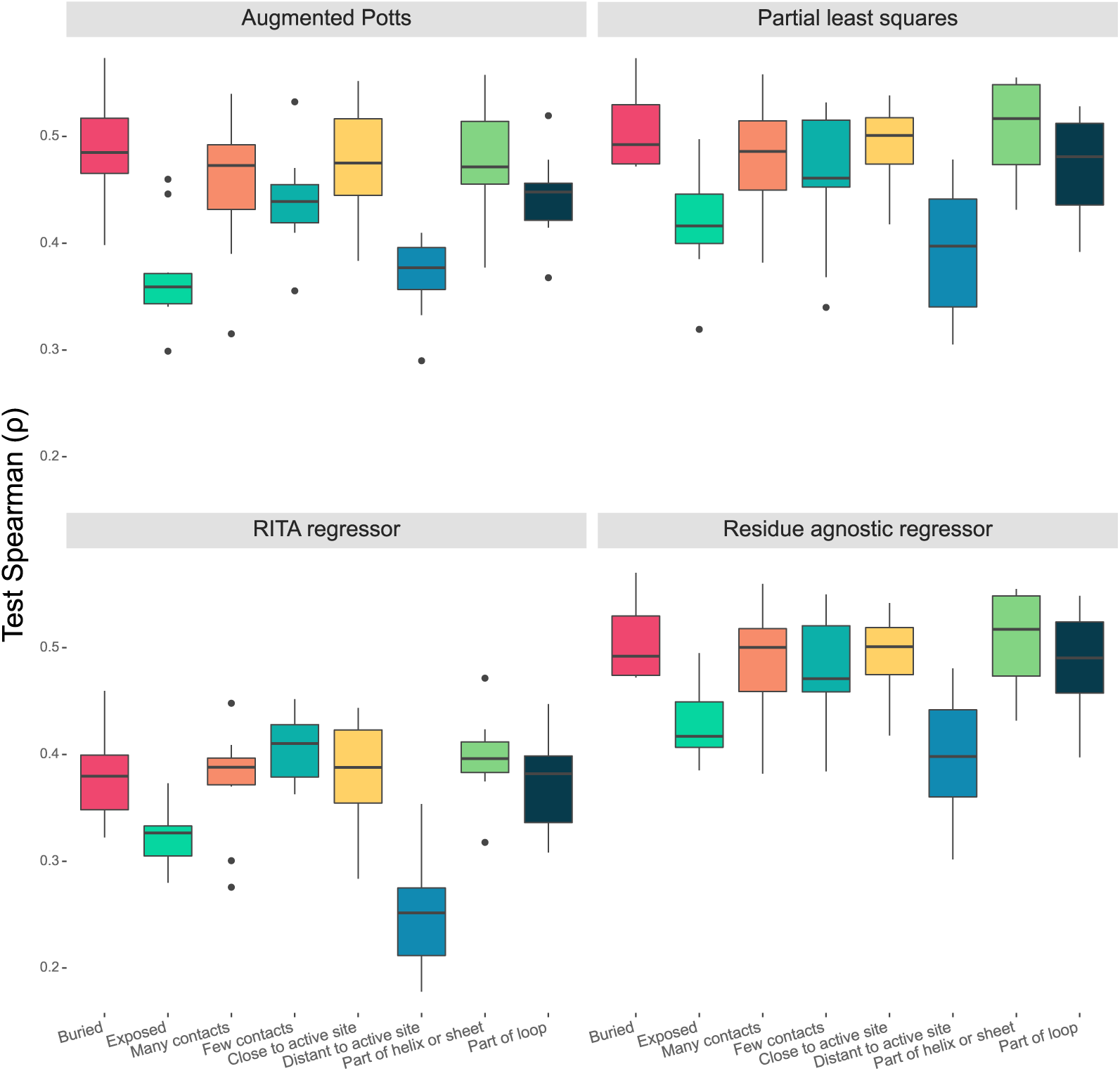
Test Spearman correlation score distributions of all models on 10 folds of the MK01_HUMAN_Brenan_2016 dataset.

**Figure S8:**
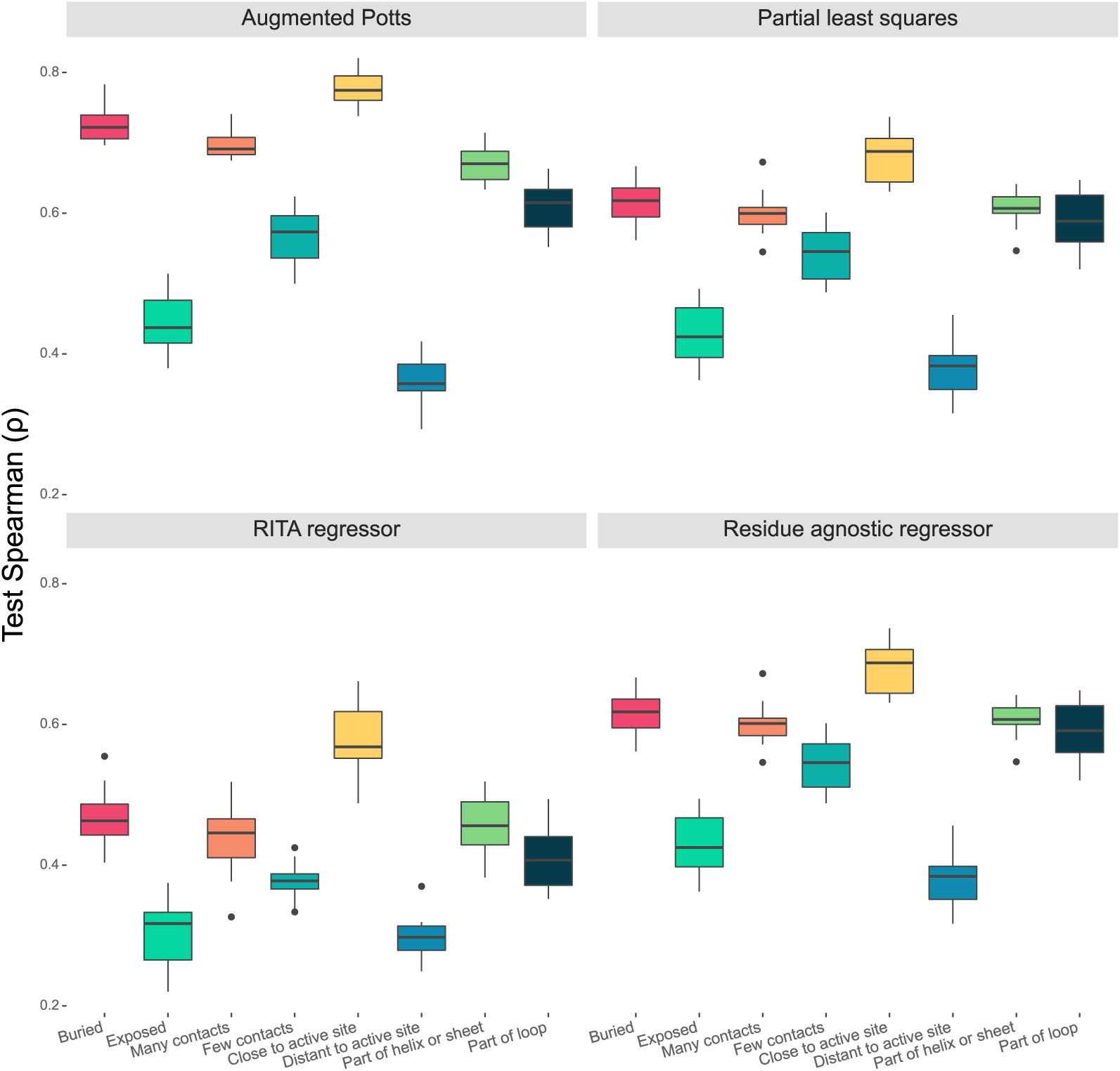
Test Spearman correlation score distributions of all models on 10 folds of the PTEN_HUMAN_Mighell_2018 dataset.

**Figure S9:**
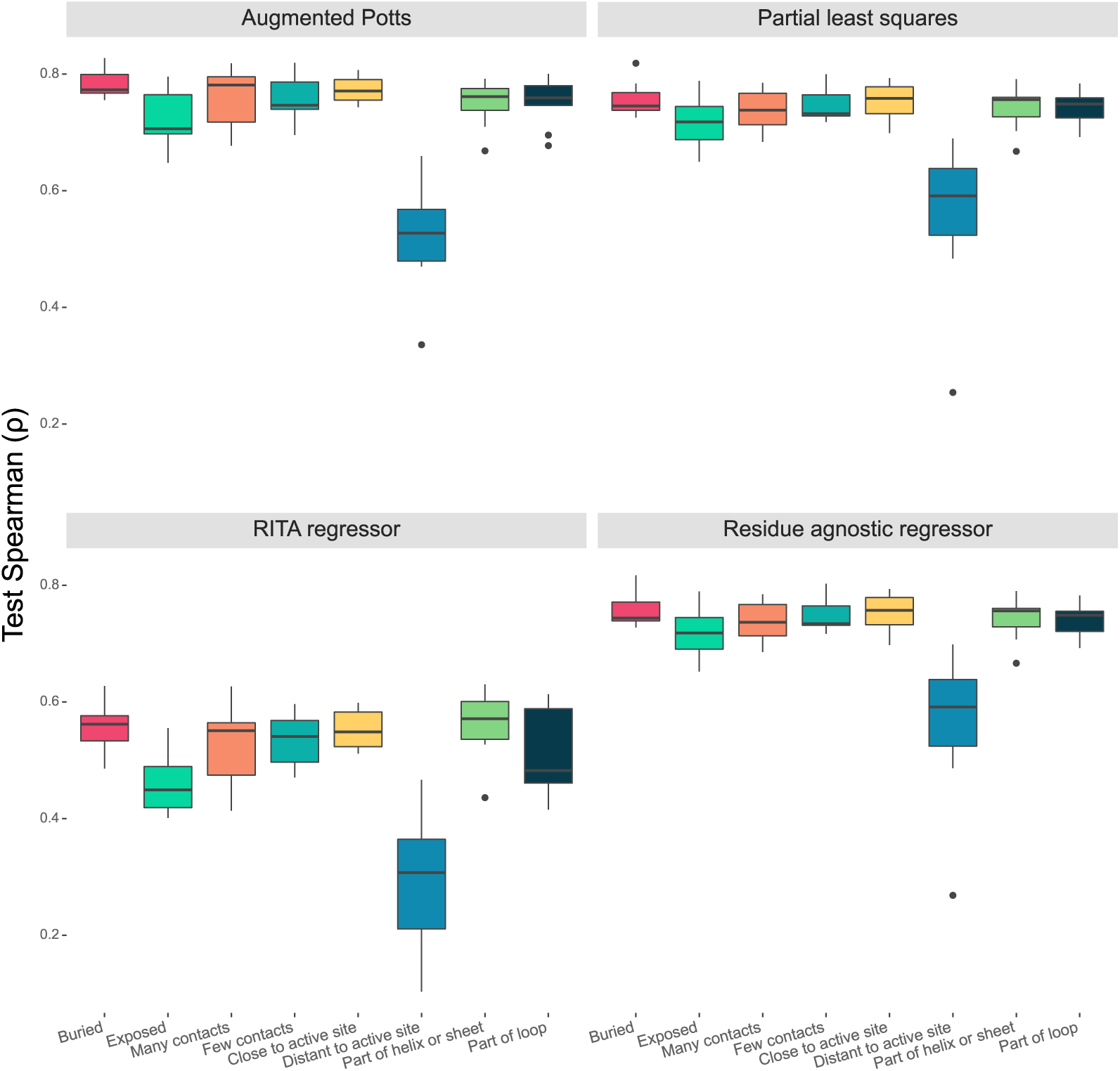
Test Spearman correlation score distributions of all models on 10 folds of the SRC_HUMAN_Ahler_2019 dataset.

**Figure S10:**
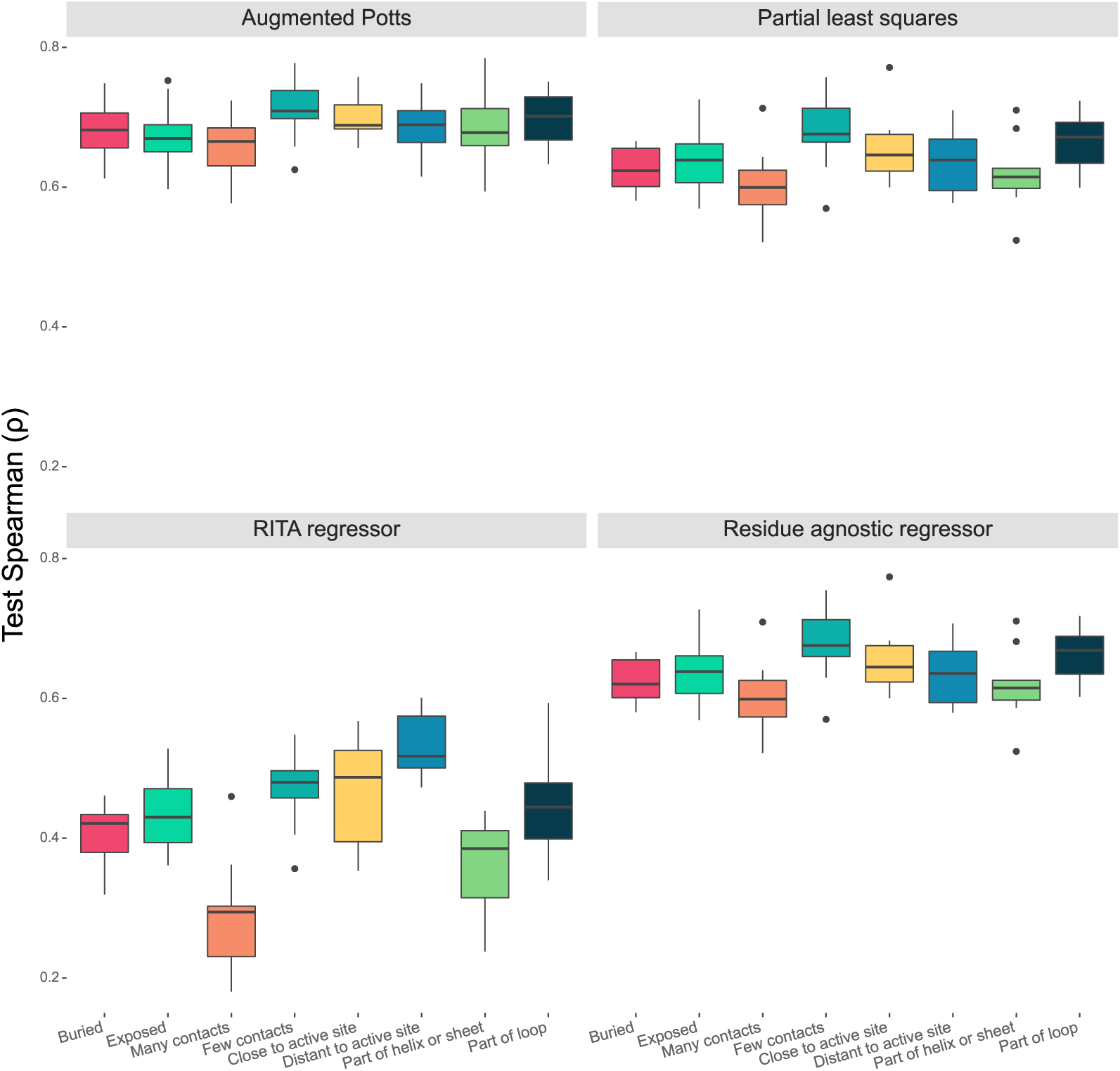
Test Spearman correlation score distributions of all models on 10 folds of the UBC9_HUMAN_Weile_2017 dataset.

**Figure S11:**
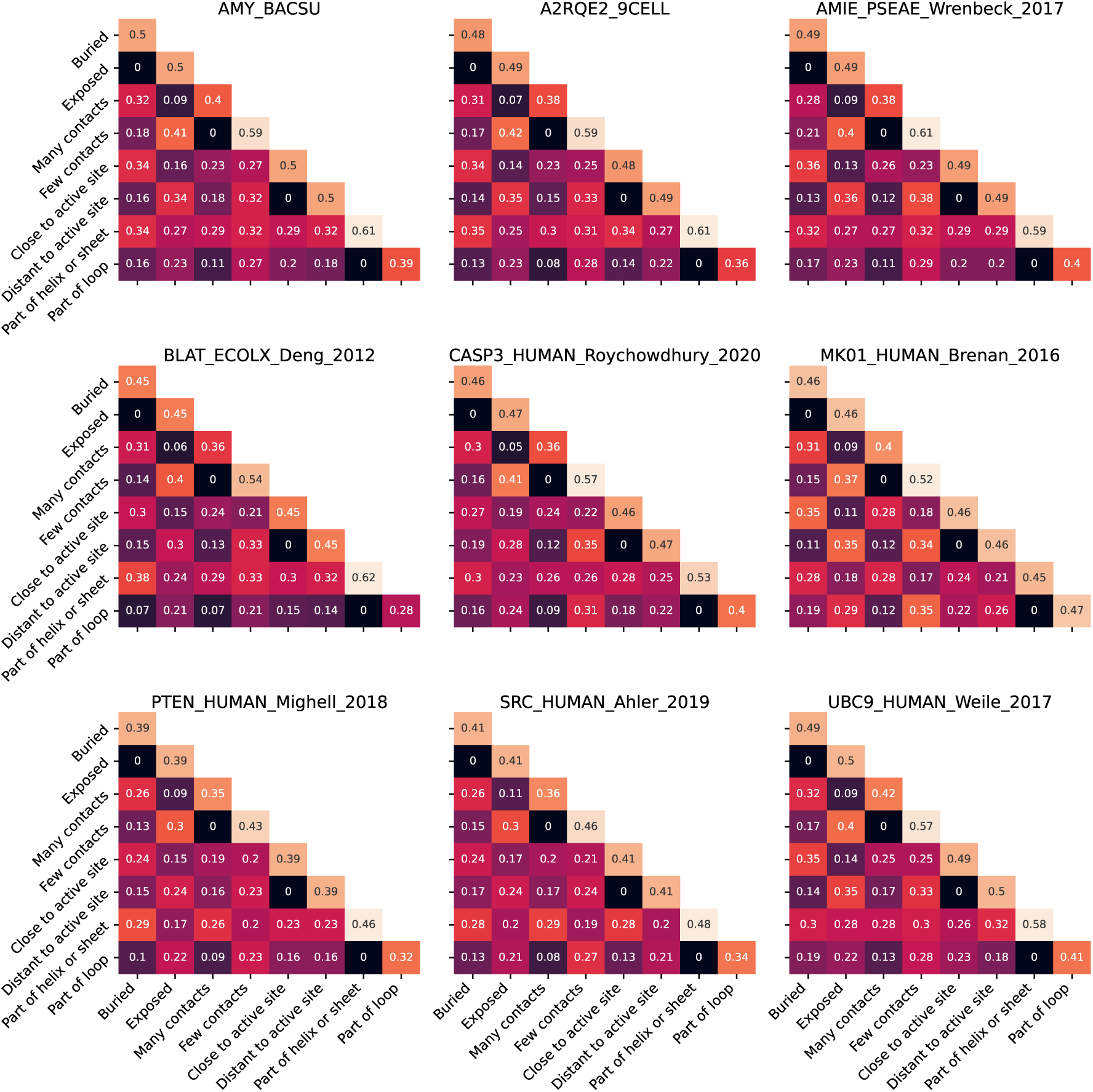
Marginal probabilities of residue membership for every pair of structural classes of all enzymes, *i.e.*, the frequencies of structurally resolved residues simultaneously being a member of structural class *C_i_* and structural class *C_j_*.

**Figure S12:**
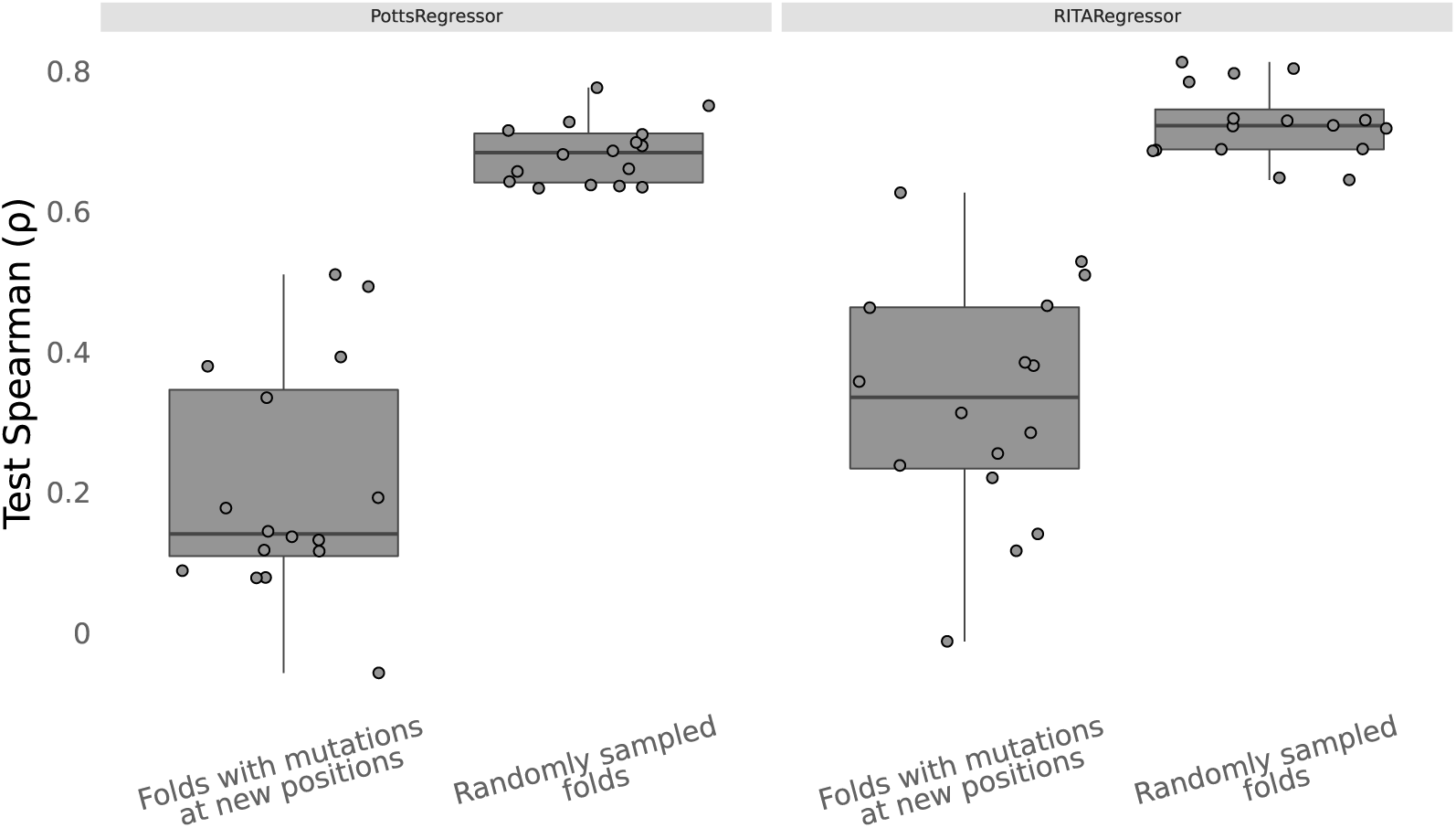
Test Spearman correlations of the augmented Potts model and RITA regressor over 16 different folds of the combinatorial variant data. Left (“Folds with mutations at novel positions”): performance on variants with mutations at positions not observed in the training data. Right (“Randomly sampled folds”): performances of models on randomly split folds.

**Table S1:**
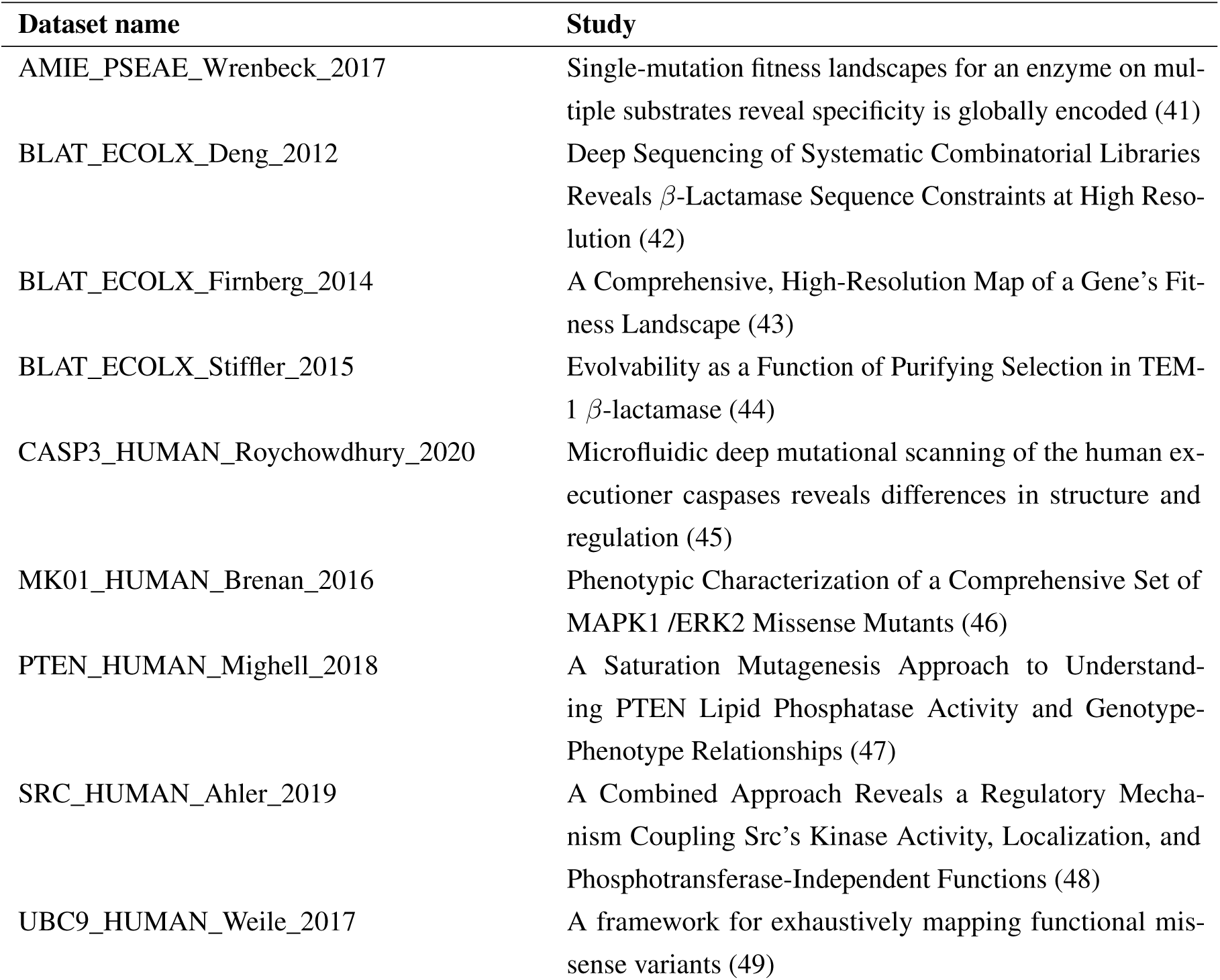
Titles of the studies corresponding to the public datasets used in our analysis.

**Figure S13:**
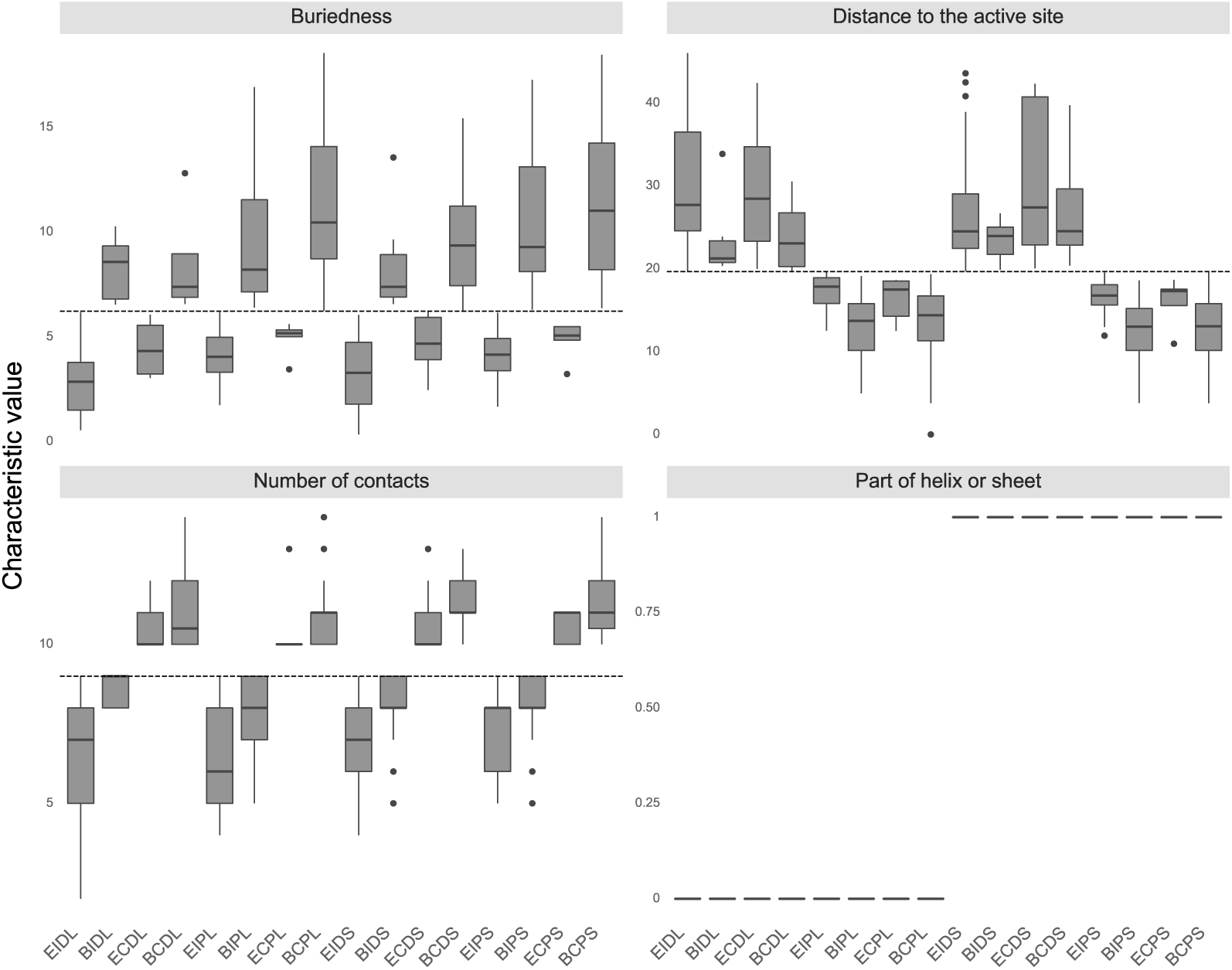
The calculated values of all structural characteristics of residues belonging to each bin. Individual bins are labeled with a four letter code specifying structural class membership. Reading from left to right, letters stand for B: Buried or E: Exposed, C: Closely connected or I: Loosely connected, P: Close to the active site or D: Distant to the active site and S: Part of helix or sheet or L: Part of loop. The median value used as threshold in assigning binary labels is visualized with the dashed line. For structural characteristics “Buriedness”, “Distance to the active site” and “Number of contacts”, bins with values below the median contain positions where residues are buried, are close to the active site and have few contact residues, respectively.

**Figure S14:**
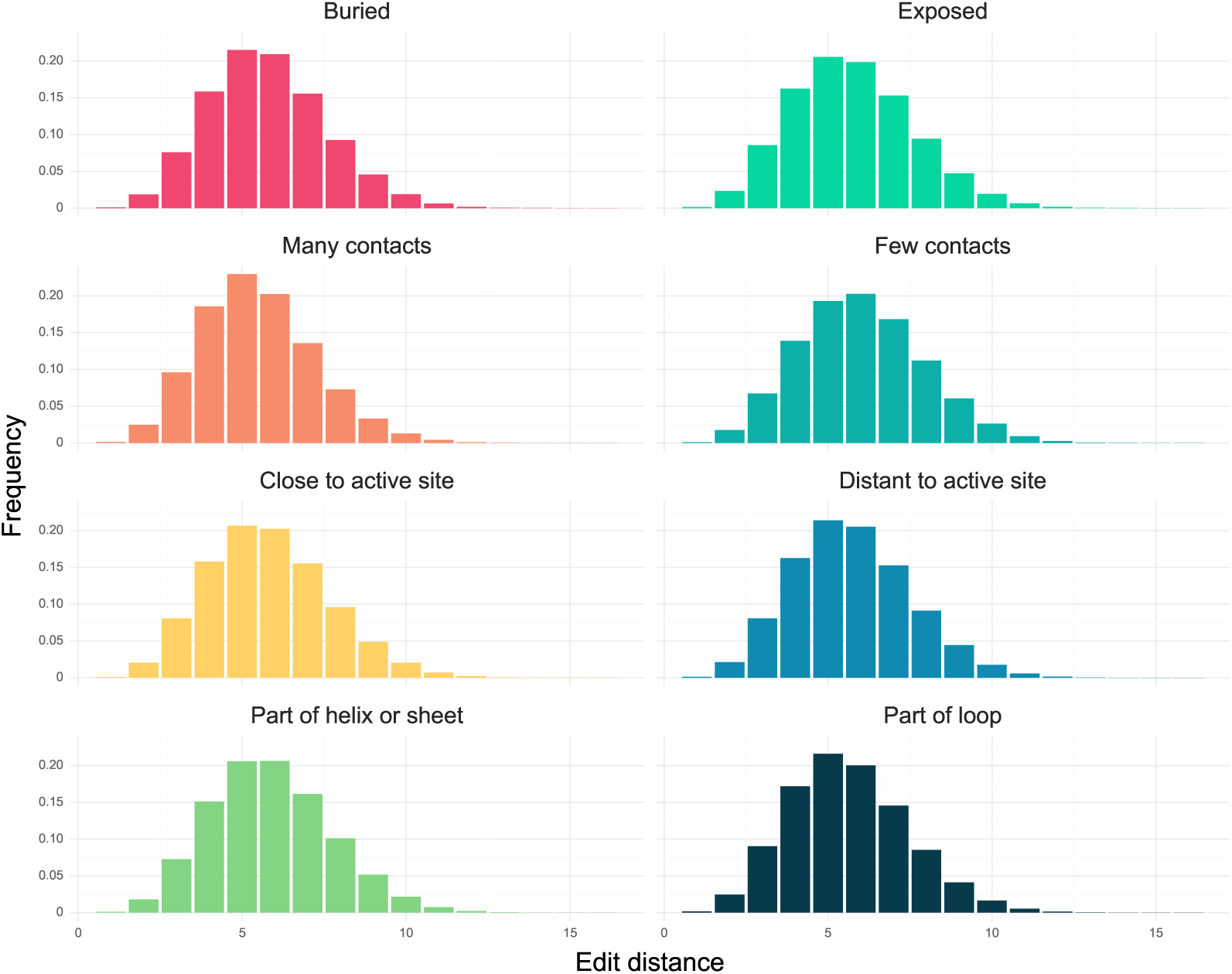
Diversity of the variants in each structural class presented as the distributions of all versus all edit distances within each structural class. Nearly identical distributions of edit distances demonstrate that the degree of variant diversity does not vary significantly among structural classes.

**Figure S15:**
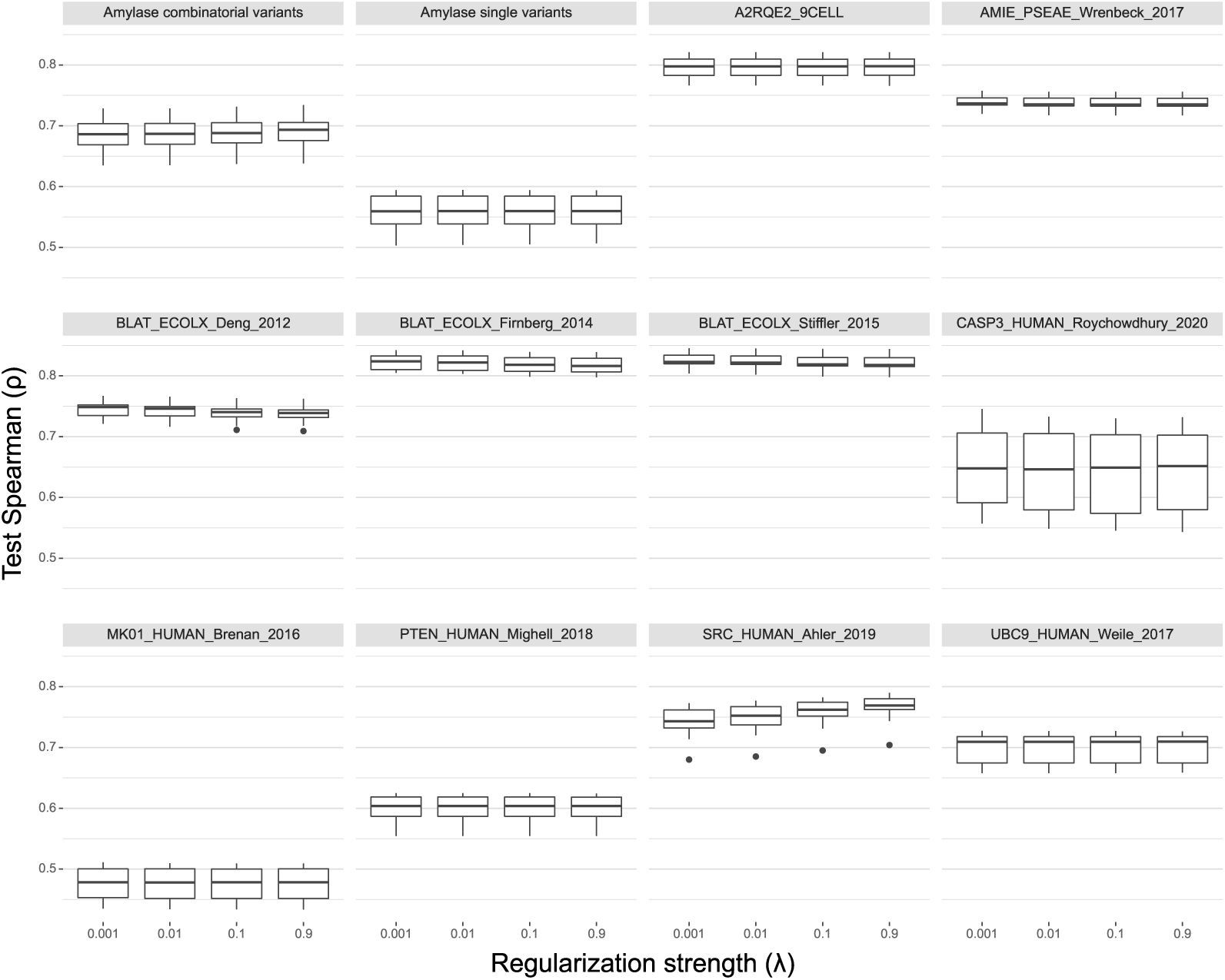
The effect of regularization strength on augmented Potts performance. Test Spearman scores are obtained from 10 cross validation of the model on each dataset in its entirety.

**Figure S16:**
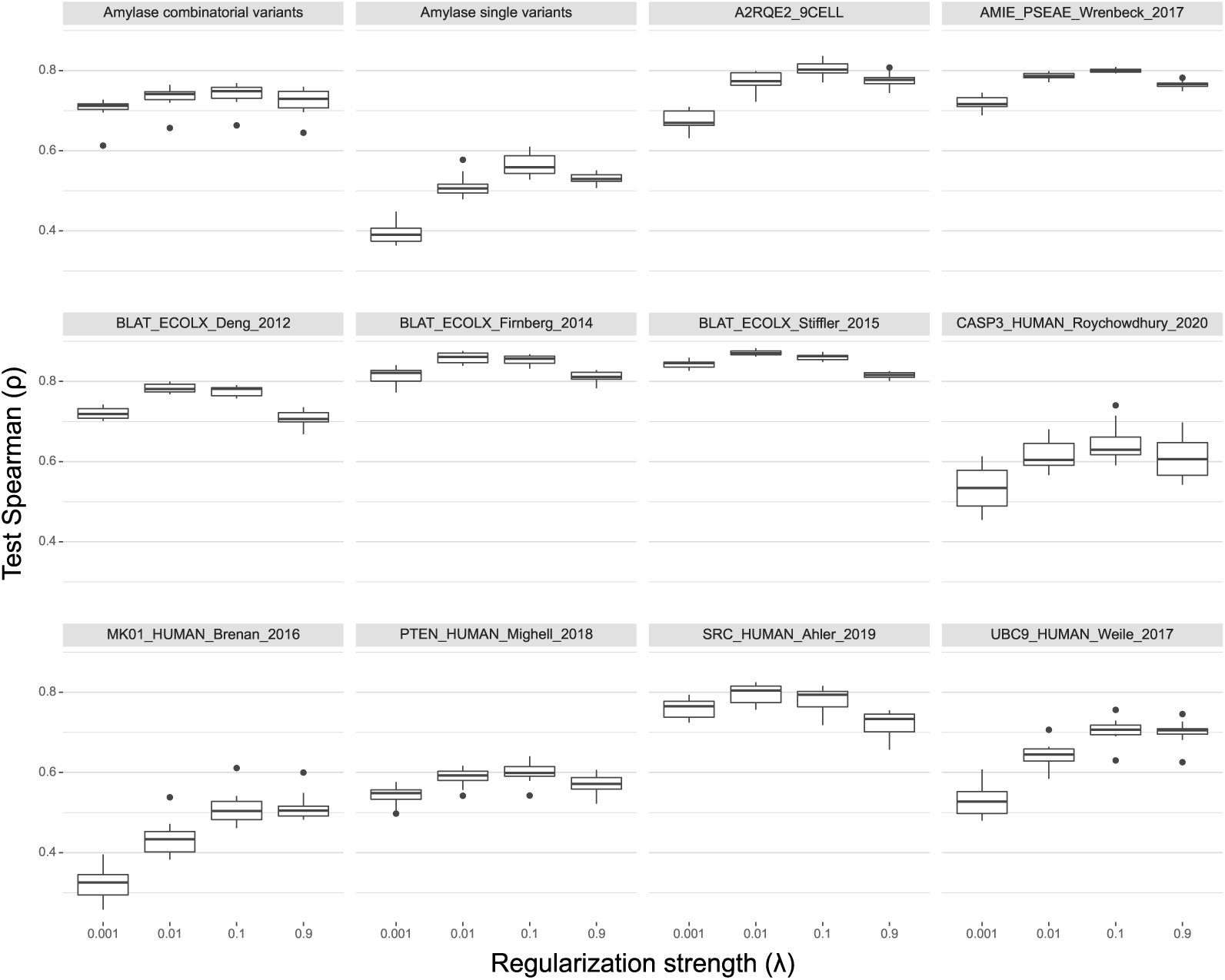
The effect of regularization strength on RITA regressor performance. Test Spearman scores are obtained from 10 cross validation of the model on each dataset in its entirety.

## Notes

https://github.com/florisvdf/mutation-predictability

### Competing Interest Statement

The authors have declared no competing interest.

### Summary of Updates

The updated manuscript contains additional experiments that were carried out using additional datasets and describes new findings.

